# Alternative splicing of a chromatin modifier alters the transcriptional regulatory programs of stem cell maintenance and neuronal differentiation

**DOI:** 10.1101/2023.09.10.556946

**Authors:** Mohammad Nazim, Chia-Ho Lin, An-Chieh Feng, Kyu-Hyeon Yeom, Mulin Li, Allison E. Daly, Xianglong Tan, Ha Vu, Jason Ernst, Michael F. Carey, Stephen T. Smale, Douglas L. Black

## Abstract

Development of embryonic stem cells (ESCs) into neurons requires intricate regulation of transcription, splicing, and translation, but how these processes interconnect is not understood. We found that polypyrimidine tract binding protein 1 (PTBP1) alters splicing of DPF2, a subunit of BAF chromatin remodeling complexes. *Dpf2* exon 7 is inhibited by PTBP1 to produce the DPF2-S isoform early in development. During neuronal differentiation, loss of PTBP1 allows exon 7 splicing, resulting in a longer DPF2-L isoform. Gene expression changes are induced by DPF2-L in ESC, and by DPF2-S in neurons. In ESC, chromatin immunoprecipitation locates DPF2-S but not DPF2-L at sites bound by pluripotency transcription factors. In neuronal progenitors, DPF2-S sites coincide with NFI protein binding, and DPF2-L sites with CTCF. DPF2-S sites show enhancer chromatin modifications, while DPF2-L sites show modifications associated with promoters. In sum, alternative splicing events during neuronal development impact chromatin organization by altering BAF complex targeting.

## INTRODUCTION

The polypyrimidine tract binding proteins PTBP1 and PTBP2 control complex programs of alternative splicing during neuronal differentiation (Boutz et al., 2007; Keppetipola et al., 2012; J. K. Vuong et al., 2016). PTBP1 is highly expressed in Embryonic Stem Cells (ESCs), where it represses inclusion of many neuronally expressed exons. PTBP1 is downregulated as ESCs differentiate into neural progenitor cells (NPCs) and neurons with a concomitant increase of PTBP2 levels (Boutz et al., 2007; Gueroussov et al., 2015; Li et al., 2014; Licatalosi et al., 2012; Linares et al., 2015; Makeyev et al., 2007). This change in regulators induces changes in the splicing of a large set of alternative exons (Boutz et al., 2007; Gueroussov et al., 2015; Li et al., 2014; J. K. Vuong et al., 2016). The resulting neuronal isoforms affect diverse functions of neuronal cells, including apoptotic potential, axonogenesis, and synaptogenesis (Lin et al., 2020; M. Zhang et al., 2019; Zheng et al., 2012). Changes in splicing can also affect transcription factors (Gabut et al., 2011; Linares et al., 2015). The PTBP1/PTBP2 splicing program thus extends into many aspects of cell biology and development.

The BRG1/BRM-associated factor (BAF) complexes are mammalian homologs of the yeast switch-sucrose non-fermentable (SWI/SNF) complex. These chromatin remodeling assemblies use ATP hydrolysis to alter chromatin conformation and modulate transcription factor (TF) access to specific genomic sites (Kadoch and Crabtree, 2015; Narayanan and Tuoc, 2014; Ronan et al., 2013; Sokpor et al., 2017). BAF complexes form polymorphic assemblies of at least 15 subunits encoded from ∼29 genes to generate diverse subunit compositions that display unique functional specificities (Kadoch and Crabtree, 2015). In early development, BAF complexes include developmental stage-specific subunits that initially control ESC pluripotency and self-renewal (Ho and Crabtree, 2010), and subsequently establish and maintain multipotent neural stem cells (NSC) and post-mitotic neurons (Kadoch and Crabtree, 2015; Sokpor et al., 2017; Son and Crabtree, 2014). As NSC-like precursors exit mitosis and acquire a neuronal fate, the BAF complex is re-organized from npBAF to nBAF. This switch involves replacing the PH10, DPF2, ACTL6A (BAF53a), and SS18 (BAF55a) subunits of npBAF with DPF1, DPF3, ACTL6B (BAF53b), and CREST (BAF55b) in nBAF, as well as changes in the levels of SMARCC1 (BAF155) and SMARCC2 (BAF170) (Lessard et al., 2007; Narayanan and Tuoc, 2014; Ronan et al., 2013; Staahl et al., 2013). Early removal of BAF53a from neural stem progenitor cells (NSPCs) results in precocious cell cycle exit and impaired cortical neurogenesis, indicating that the timely switch from npBAF to nBAF is critical for proper cortical development (Braun et al., 2021).

The BAF subunit DPF2 is one of a family of four paralogous genes: DPF1 (BAF45b), DPF2 (BAF45d), DPF3 (BAF45c), and PH10 (BAF45a). These BAF45 paralogs each contain two C-terminal plant homeodomain (PHD) finger domains that target the BAF complex to genomic loci bearing specific histone marks and alter the role of BAF in different developmental lineages (Kadoch and Crabtree, 2015; Kulikova et al., 2013; Vasileiou et al., 2018). PH10 plays a critical role in maintenance of hematopoietic stem cells (Krasteva et al., 2017), DPF3 in heart and muscle development (Lange et al., 2008), and both DPF1 and DPF3 in self-renewal and differentiation of NPCs (Kadoch and Crabtree, 2015). DPF2 is more broadly expressed than the other BAF45 paralogs, and has been implicated in diverse functions including programmed cell death (apoptosis) after IL-3 deprivation in myeloid cells (Gabig et al., 1994), interaction with pluripotency transcription factors in stem cells (Pardo et al., 2010; van den Berg et al., 2010), and expression of the T-Box Transcription Factor 3 (Tbx3) during mesendodermal differentiation (W. Zhang et al., 2019). The relationship of the neuronal BAF transcriptional regulatory program to the posttranscriptional program controlled by the PTB proteins is not well understood.

We found that an additional BAF subunit is expressed in brain, where *Dpf2* transcripts are alternatively spliced to include exon 7. This creates a longer DPF2-L protein isoform that replaces the canonical short DPF2-S isoform in neurons. We show that DPF2 exon 7 is repressed by PTBP1 in early development but becomes included in the *Dpf2* mRNA as PTBP1 is depleted during neuronal maturation. Expression of DPF2-S or DPF2-L in ESCs, NPCs, and glutamatergic neurons drives different gene expression programs, and yields chromatin binding profiles that are cell type specific. In ESCs, DPF2-S preferentially bound to chromatin sites associated with pluripotency transcription factors, while in NPC DPF2-L sites overlapped with binding sites for the factor CTCF. DPF2-S and DPF2-L binding sites were further distinguished by their chromatin state enrichments, with DPF2-S sites associated with enhancer chromatin states, and DPF2-L sites associated with states indicative of transcription start sites and promoters. Overall, we uncover a new interplay between the posttranscriptional splicing regulatory program directed by the PTB proteins during neuronal development and chromatin level regulation mediated by neuronal BAF complexes.

## RESULTS

### *Dpf2* exon 7 is spliced in a tissue- and developmental stage-specific manner

Studies of alternative splicing in mouse brain revealed that *Dpf2* transcripts contained an additional exon (exon 7), not seen in the canonical *Dpf2* mRNAs of other tissues **(Figure 1A)**. The new exon 7 is conserved across vertebrate species and its inclusion inserts a new 14 amino acid peptide into the C2H2-type zinc finger domain of DPF2 protein **(Figures S1A-B)**. Inclusion of exon 7 in DPF2-L splits the two critical Cysteine residues, disrupting the consensus C2H2-type zinc finger motif, and introducing several basic amino acids (three Lysine residues and one Arginine residue) **(Figure S1B)**. C2H2-type zinc finger domains frequently engage in DNA, RNA, and protein interactions (Brayer and Segal, 2008) and we hypothesized that introduction of the new peptide will alter the zinc finger’s interactions and DPF2 function.

**Figure 1.**
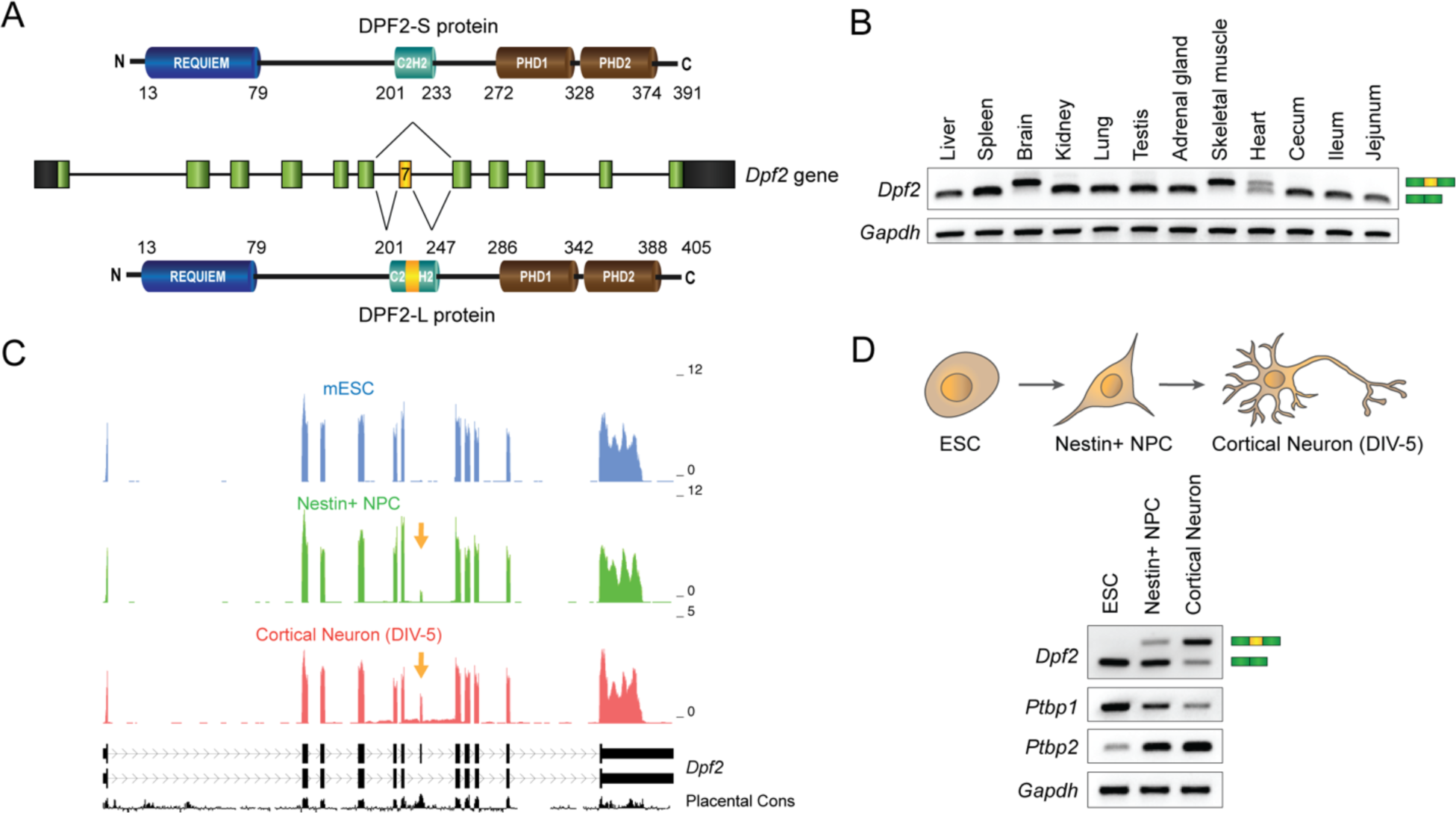
*Dpf2* exon 7 is alternatively spliced in a tissue- and developmental stage-specific manner. **(A)** Schematic of *Dpf2* gene and alternatively spliced variants of DPF2 protein. Exon 7 is alternatively spliced to either generate the canonical DPF2-S protein, or a longer DPF2-L with a 14 amino acid insert in the C2H2-type zinc finger domain. The numbers denote amino acid numbers. REQUIEM, C2H2, PHD1, and PHD2 are known functional domains of DPF2. PHD, plant homeodomain. **(B)** RT-PCR assay monitoring *Dpf2* exon 7 alternative splicing across different mouse tissues. *Gapdh* is shown as a loading control. **(C)** Genome browser tracks of the *Dpf2* locus from mouse ESCs, Nestin^+^ NPCs isolated from brain, and mouse cortical neurons cultured in vitro for 5 days (DIV-5). Inclusion of *Dpf2* exon 7 is indicated by yellow arrows. *Dpf2* gene structure and vertebrate conservation is shown below. **(D)** RT-PCR assays showing *Dpf2* exon 7 alternative splicing in ESCs, NPCs, and cortical neurons (DIV-5). Expression levels of *Ptbp1* and *Ptbp2* are shown below. *Gapdh* is shown as a control. See also **Figure S1**.

RT-PCR assays of RNA from different mouse tissues indicated that exon 7 is included in nearly 100% of the *Dpf2* mRNA in brain and muscle, and ∼50% in heart, but is almost entirely skipped in all other tissues examined **(Figure 1B)**. RT-PCR analysis in mouse cortex at developmental times from E15 to P60 showed that exon 7 is increasingly included as neuronal development progresses **(Figure S1C)**. Published RNA-seq data (Yeom et al., 2021) showed that *Dpf2* exon 7 is entirely skipped in mouse embryonic stem cells (ESCs), weakly included in neuronal progenitor cells (NPCs), and predominantly included in cortical neurons differentiated for 5 days *in vitro* **(Figure 1C)**. These observations were validated by RT-PCR analysis in the same cells, which showed that exon 7 inclusion continues to increase with continued neuronal maturation beyond DIV-5 **(Figure 1D)**. Both in cell culture and in brain, the induction of exon 7 splicing coincides with the decline in PTBP1 expression and the concomitant increase in PTBP2 expression, implicating PTBP1 as a candidate regulator of exon 7 splicing **(Figure 1D)**. The switch in DPF2 exon 7 splicing from nearly complete skipping in ESCs to efficient inclusion in differentiated neurons is similar to other targets of PTBP1 (Linares et al., 2015; Zheng et al., 2012).

### The RNA binding protein PTBP1 and a weak 5’ splice site together regulate the alternative splicing of *Dpf2* exon 7

Five blocks of pyrimidine-rich sequence in the introns upstream and downstream of exon 7 are potential binding sites for PTBP1 **(Figures 2C, S2C)** (Han et al., 2014; Keppetipola et al., 2012; Llorian et al., 2010; Markovtsov et al., 2000; Spellman et al., 2007; J. K. Vuong et al., 2016). Individual-nucleotide resolution cross-linking and immunoprecipitation (iCLIP-seq) data from mouse ESCs confirmed PTBP1 binding to the exon 7 region, with one crosslinking site within the 3’ splice site, and another cluster of crosslinking sites ∼40 nt downstream of the 5’ splice site (SS) (Linares et al., 2015) **(Figure 2A)**. This arrangement of flanking binding sites is characteristic of exons repressed by PTBP1 (Amir-Ahmady et al., 2005; Ye et al., 2023). To further examine the sequences regulating exon 7, we constructed a minigene reporter containing the sequence from *Dpf2* exon 6 to exon 8 along with introns 6 and 7. We performed RNAi knockdown of PTBP1 and its paralog PTBP2 in mouse ESCs followed by transient transfection of the DPF2 minigene. We found that depletion of PTBP1 alone induced exon 7 inclusion from 0 to ∼60% in ESC **(Figure 2B)**. As expected, PTBP1 depletion induced expression of its paralog PTBP2, but simultaneous depletion of PTBP1 and PTBP2 had only a small additional effect over PTBP1 depletion alone **(Figure 2B)**. Importantly, exon 7 from the endogenous DPF2 locus is also strongly included upon depletion of PTBP1 either alone or simultaneously with PTBP2. We obtained similar results with a different set of PTBP1 and PTBP2 siRNAs, indicating that the splicing changes were not off-target effects of the knockdowns **(Figures S2A-B)**. Altogether, the data indicate that splicing of DPF2 exon 7 is primarily regulated by PTBP1, consistent with its switch during development when PTBP1 is downregulated.

**Figure 2.**
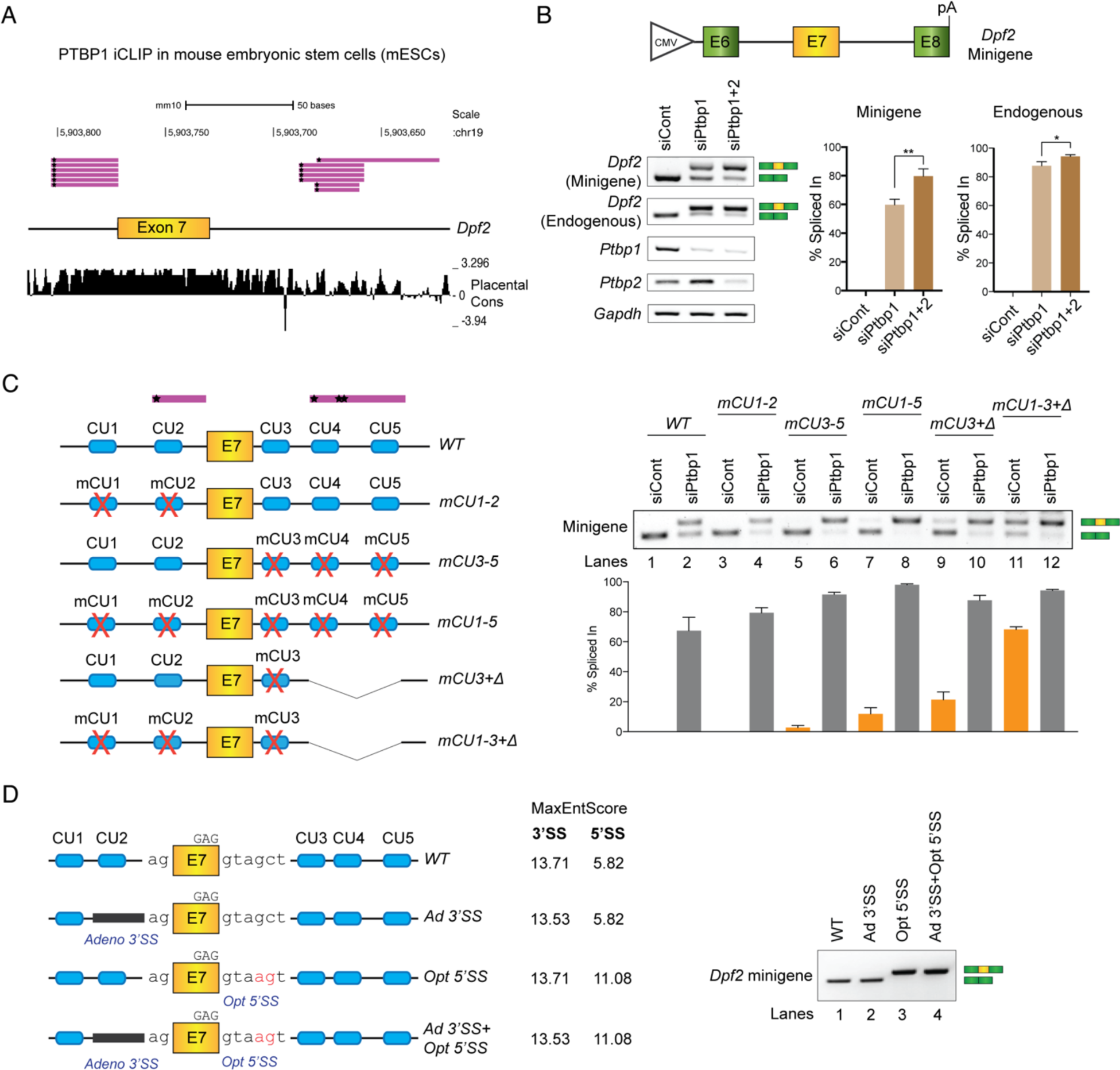
The RNA binding protein PTBP1 and a weak 5’ splice site together regulate the alternative splicing of *Dpf2* exon 7. **(A)** Genome browser tracks showing PTBP1 iCLIP-tags in the flanking intronic regions of *Dpf2* exon 7 in mouse ESCs. Individual iCLIP clusters are shown as purple lines, where the ‘star’ mark denotes the cross-link site for each tag. **(B)** Diagram of *Dpf2* minigene spanning E6-E8 region of *Dpf2* gene. RT-PCR assay of exon 7 alternative splicing in *Dpf2* minigene and endogenous *Dpf2* transcripts after siRNA-mediated knockdown of *Ptbp1* and *Ptbp2*. Expression of *Ptbp1*, *Ptbp2*, and *Gapdh* (loading control) are also shown. Bar graphs (right) show quantification of RT-PCR results calculated as exon 7 percent spliced in (PSI) for the minigene and endogenous *Dpf2* transcripts. Each bar represents the mean value (+/-SD) of three independent biological replicates. *: p < 0.05; **: p < 0.01 (Student’s t-test) **(C)** Schematics of *Dpf2* minigenes containing wild-type or mutated CU-rich elements flanking exon 7. The red ‘X’ denotes mutated CU rich elements, and delta denotes deletion of a 67-nt pyrimidine-rich region. Positions of the PTBP1 iCLIP clusters are indicated by purple bars above. The right panel shows quantification of RT-PCR data monitoring exon 7 splicing in the minigene transcripts. Each bar represents the mean value (+/-SD) of three independent biological replicates. **(D)** Schematic of *Dpf2* minigenes containing wild-type or mutated 5’ and 3’ splice sites. Corresponding MaxEntScan scores (Yeo and Burge, 2003) of 5’ and 3’ splice sites are shown on the right. The right panel shows RT-PCR assays of exon 7 splicing in minigene constructs containing wild-type or mutated 5’ and 3’ splice sites. See also **Figure S2**.

To assess the cis-regulatory elements affecting exon 7 splicing, we constructed minigenes carrying mutations in the CU-rich motifs. Mutation of the two upstream elements together, the three downstream elements together, or all five CU-rich elements together only marginally affected exon 7 splicing **(Figure 2C**, lanes 3, 5, 7), suggesting that there are additional repressive regulatory elements. However, after PTBP1 depletion, exon 7 was more strongly included in these mutants than in the wild-type minigene, indicating that the CU-rich elements are contributing to exon 7 repression **(Figure 2C**, lanes 2, 4, 6, 8). To inactivate additional elements, we deleted the intronic region containing the downstream PTBP1 iCLIP cluster, along with either mutating the remaining downstream CU-rich element or mutating all the remaining upstream and downstream elements. With only the upstream elements remaining, exon 7 splicing was enhanced, showing partial inclusion even in the presence of PTBP1 (mCU3+Δ; **Figure 2C**, lane 9). When all the CU element mutations were combined with the deletion, exon 7 became predominantly included in the mRNA even in the presence of PTBP1 (mCU1-3+Δ; **Figure 2C**, lane 11). These data indicate that DPF2 exon 7 is highly sensitive to the presence of PTBP1, which is binding to distributed sites in its flanking introns.

To assess other contributions to exon 7 regulation, we examined the strength of its 5’ and 3’ splice sites using the MaxEntScan algorithm (Yeo and Burge, 2003). The 5’ splice site at the exon 7-intron 7 junction is suboptimal with a MaxEntScan score of 5.82, while the 3’ splice site at the intron 6-exon 7 junction is strong (MaxEntScan score = 13.71). We found that mutating the 5’ splice site to the consensus sequence (MaxEntScan score = 11.08) led to nearly complete exon 7 inclusion in mouse ESCs, even with the CU elements intact and in the presence of PTBP1 **(Figure 2D**, lane 3, **Figure S2D)**. Optimizing the 5’ splice site can thus override repression of the exon by PTBP1. Because the iCLIP tag and one of the CU-rich elements upstream of exon 7 overlap with the polypyrimidine track of the 3’ splice site, we replaced this site with a strong polyU-rich 3’ splice site derived from an intron of the adenovirus major late transcription unit that is not repressed by PTBP1 (Chan and Black, 1997). This new 3’ splice site did not alter exon 7 splicing on its own **(Figure 2D**, lane 2), and conversion of both splice sites together did not elicit additional effects over the optimal 5’ splice site alone **(Figure 2D**, lane 4). In sum, DPF2 exon 7 is part of the large program of splicing changes controlled by PTBP1. The strong regulation of this exon during development and its conservation across vertebrate species imply an important function and we next wanted to assess how inclusion of exon 7 affected DPF2 activity.

### Neuronal differentiation of mouse ESCs expressing either DPF2-S or DPF2-L

To examine the functions of DPF2-S and DPF2-L at different stages of development, we modified the DPF2 locus in mouse ESC cells using CRISPR/Cas9-mediated genome editing. We used a pair of guide RNAs (gRNAs) targeting intronic sites to delete exon 7 (E7-KO) and adjacent intron sequences to create cells that only express DPF2-S **(Figures 3A, S2E-F)**. We used the same gRNAs with a homology directed repair construct carrying a modified exon 7 (E7-KI) with strong 5’ and 3’ splice sites and no nearby CU-rich elements to induce constitutive splicing of exon 7 **(Figures 3A, S2E, S2G)**. We then screened for mouse ESC clones homozygous for the two alleles **(Figure 3B)**. RT-PCR and western blot assays confirmed that the wild-type and E7-KO cells expressed the DPF2-S isoform and the E7-KI cells exclusively express the DPF2-L isoform **(Figure 3C)**. We then expanded three clones of each genotype.

**Figure 3.**
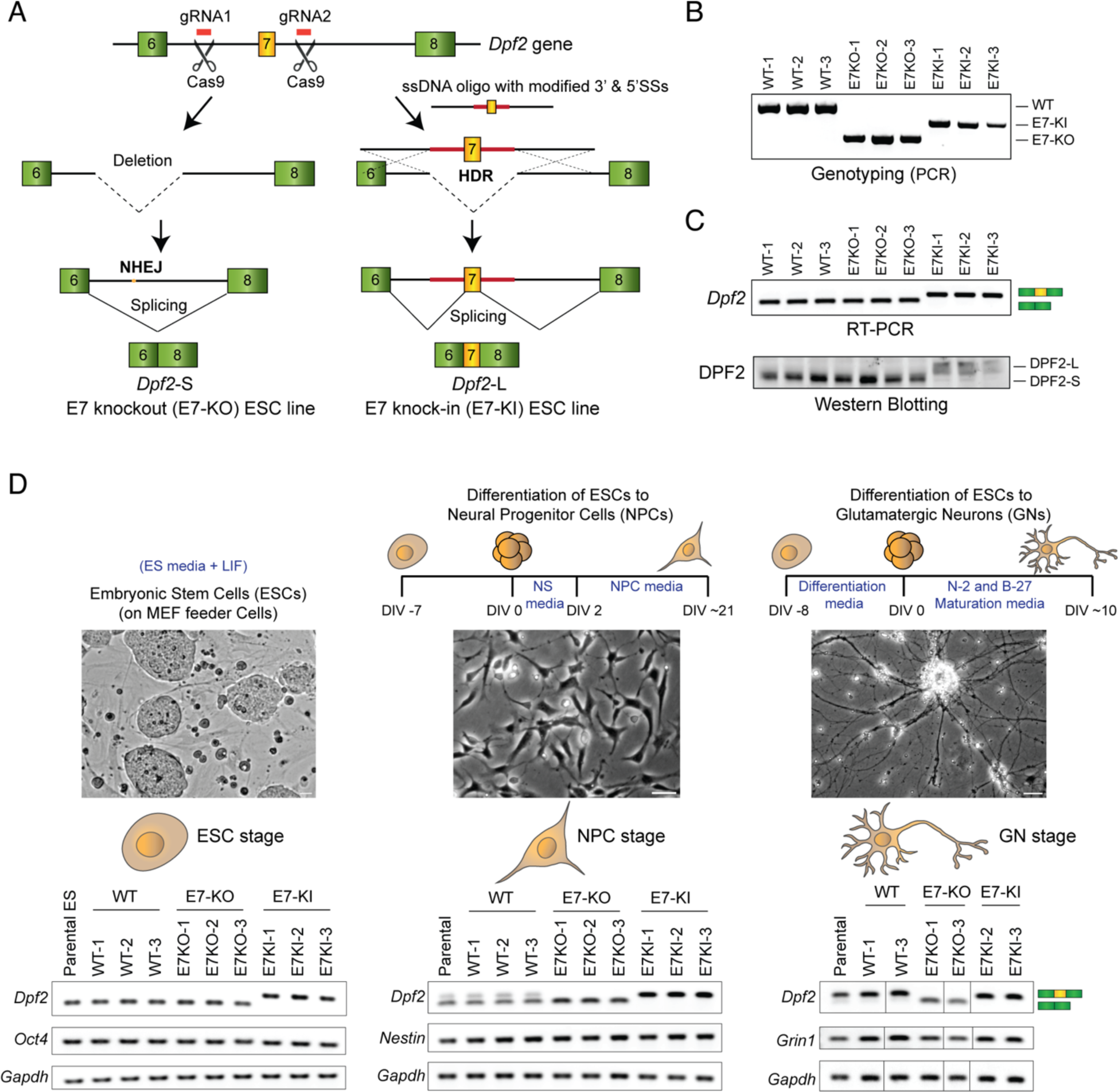
Forced expression of either DPF2-S or DPF2-L in genome-edited mouse ESCs and their differentiation to NPCs and glutamatergic neurons. **(A)** Schematic of the CRISPR/Cas9-mediated genome editing strategy for generating cells that exclusively express either DPF2-S, or DPF2-L. Two guide RNAs targeted sites in intron 6 and intron 7 to delete exon 7 with its flanking intronic sequences, producing a gene that only expresses DPF2-S isoform. In a second ESC line, these two guide RNAs were introduced with a repair ssDNA template containing optimal 5’ and 3’ splice sites flanking exon 7 and lacking CU-rich elements. This creates a gene that only produces the DPF2-L isoform with exon 7 included. **(B)** Genotyping of individual CRISPR/Cas9 genome-edited ESC clones carrying homozygous wild-type, exon 7 deleted (E7-KO), and modified exon 7 knock-in (E7-KI) alleles. **(C)** RT-PCR assay of exon 7 splicing and immunoblot of the DPF2 protein isoforms in wild-type and genome edited ESC clones. **(D)** Schematic showing the strategy of differentiating ESCs to NPCs cultured for 21 days in vitro (DIV-21), or to glutamatergic neurons (GNs) cultured for 10 days in vitro (DIV-10), with representative cell images. The bottom panel shows RT-PCR assays of exon 7 splicing in wild-type and genome-edited ESCs, NPCs, and GNs. Expression levels of *Oct4*, *Nestin*, and *Grin1* are shown as marker genes for each developmental stage. Expression levels of *Gapdh* are shown as a control. See also **Figure S2**.

The ESC clones were aggregated as embryoid bodies (EBs) in differentiation medium and then differentiated into neural progenitor cells (NPCs) by culturing them in NPC medium for 21 days (DIV-21), when they expressed the NPC marker *Nestin* and had developed NPC morphology with short membrane processes. At DIV-21, wild-type NPCs showed partial inclusion of DPF2 exon 7, while the E7-KO cells expressed only the DPF2-S variant, and the E7-KI cells expressed only the DPF2-L variant **(Figure 3D)**. We also differentiated the ESC clones from EBs into post-mitotic glutamatergic neurons (GNs) in maturation medium until DIV-10 when they were expressing the neuronal marker *Grin1* and had developed the extended thin processes of differentiated neurons. One clone of each of the three genotypes did not differentiate well into glutamatergic neurons and was discarded. The remaining clones all developed normally. Overall, we did not observe differences between the three genotypes in the efficiency of NPC or GN differentiation. As expected, the wild-type GNs almost exclusively expressed DPF2-L by DIV-10, as did the E7-KI cells. In contrast, the E7-KO cells exclusively expressed DPF2-S in the now differentiated neurons **(Figure 3D)**. Collectively, these cell lines allow us to evaluate DPF2-S and DPF2-L function during neuronal development.

### The alternative DPF2 isoforms regulate distinct programs of gene expression during neuronal differentiation

To determine how alternative DPF2 variants control the transcriptome during development, we performed RNA-seq profiling. We analyzed the two clones of each genotype to yield biological duplicates in ESCs, NPCs, and neurons **(Figure S3A)**. RNA-seq reads from each cell clone at each developmental stage were mapped to the mouse genome (NCBI38/mm10) with RPKM values determined in annotated genes (Mortazavi et al., 2008). To provide the clearest comparison of the two DPF2 isoforms, average RPKM values from the E7-KO cells (only express DPF2-S) were compared to RPKM values from the E7-KI cells (only express DPF2-L). Applying cutoffs of >1.5-fold change in expression between these two cells with a *P* value below 0.05 (FDR < 0.1), we used DE-seq2 to identify 372 genes more highly expressed in the E7-KO ESCs and 509 genes more highly expressed in the E7-KI cells **(Figures 4A, S3B)**.

**Figure 4.**
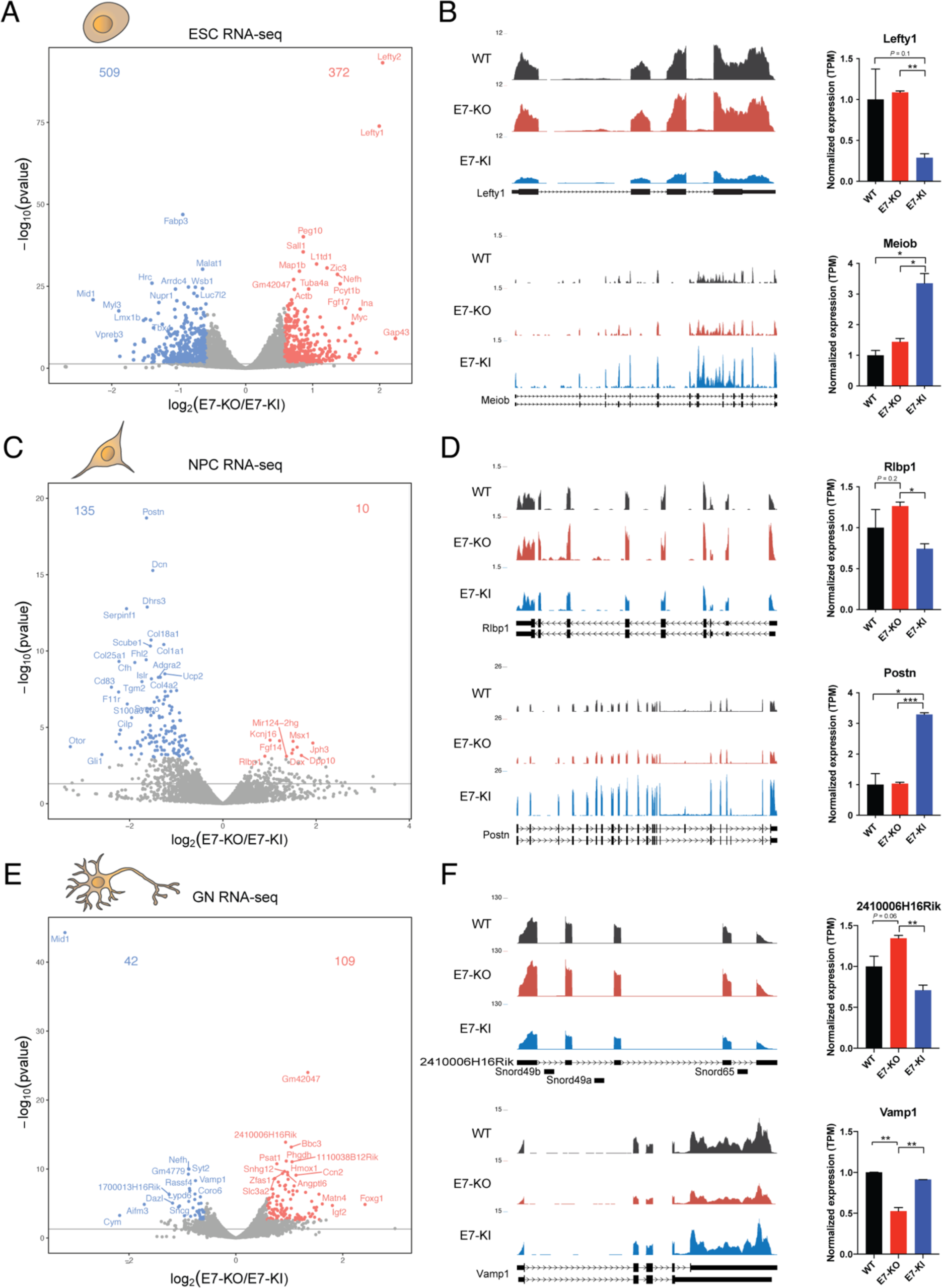
Alternative DPF2 isoforms regulate distinct programs of gene expression during neuronal differentiation of ESCs. **(A, C, E)** Volcano plots showing differentially expressed genes (DEGs) detected by RNA-seq in mouse ESCs (A), ESC-derived NPCs (C), and ESC-derived GNs (E) expressing either DPF2-S or DPF2-L. Significant DEGs were filtered by 1.5-fold changes with a *P*-value below 0.05 (FDR < 0.1) between E7-KO and E7-KI and labeled in red and blue dots. **(B, D, F)** Genome browser tracks show aligned RNA-seq reads of individual DEGs in wild-type (WT), DPF2-S expressing (E7-KO), and DPF2-L expressing (E7-KI) ESCs (B), NPCs (D), and GNs (F). The right panel shows bar charts of normalized expression (TPM) of individual DEGs in these cell lines. See also **Figure S3**.

Gene Ontology (GO) analysis revealed that the genes upregulated in E7-KO ESCs relative to the E7-KI were significantly enriched for terms such as tissue morphogenesis, epithelial cell differentiation, cell morphogenesis involved in differentiation, mechanisms associated with pluripotency, embryonic morphogenesis, and others. The GO terms for the genes upregulated in E7-KI ESCs were not as strongly enriched as the E7-KO ESCs, and included terms such as meiotic cell cycle, and chloride transport **(Figure S3B)**. Interestingly, among the genes most strongly upregulated in the E7-KO ESCs over the E7-KI ESCs were the stem cell identity genes *Lefty1* and *Lefty2*, indicating that DPF2-S isoform affects maintenance of pluripotency **(Figures 4A-B)**. Other pluripotency-associated genes such as *Myc, Tcf7l1, Tdgf1, Wt1, Bmp4, Zic2/3, Otx2, Nodal, Lef1,* and *Tcf15* were also upregulated in E7-KO ESCs expressing the DPF2-S isoform.

Using similar expression cutoffs at the NPC stage, we found only 10 genes to be higher in the E7-KO NPCs and 135 genes to be higher in E7-KI NPCs **(Figures 4C-D, S3C)**. The most enriched GO term for genes upregulated in E7-KI NPCs was extracellular matrix organization, which included genes such as *Loxl1/2*, *Adamts1/2*, *Ddr2*, *Cyp1b1*, *Fkbp10*, *Emilin1*, *Mmp2*, *Fn1*, *Postn*, and multiple collagen genes *Col1a1*, *Col8a1*, etc. In differentiated neurons, we found 109 genes to be more highly expressed in E7-KO GNs and 42 genes to be higher in E7-KI GNs **(Figures 4E-F, S3D)**. Notably, neuron-specific genes such as *Vamp1*, *Syt2*, and *Nefh* were among the genes strongly upregulated in E7-KI GNs. Other upregulated neuron-specific genes associated with synaptic transmission and neuroactive ligand-receptor interaction were *Rph3a, Sncg, Glra3, Lynx1, Hapln4,* and *Chrm2*, suggesting that DPF2-L isoform modulates a subset of neuronal genes. Based on fewer genes, GO analyses of the differentially expressed genes in NPC’s and GN yielded fewer enrichments **(Figures S3C-D)**.

These data indicate that the two DPF2 isoforms are functioning differently at different stages of development, with DPF2-S affecting genes contributing to pluripotency in stem cells, and DPF2-L altering neuron-specific genes after differentiation. The two DPF2 isoforms do not seem to determine the on/off state of these genes but instead modulate their overall expression. We next wanted to examine whether these different activities of the two isoforms were reflected in their targeting to chromatin.

### DPF2-S preferentially targets sequences bound by stem cell pluripotency factors in ESCs

To identify genes bound by the two alternative DPF2 isoforms, we performed chromatin immunoprecipitation assays (ChIP-seq) in mouse ESCs. Using an antibody that efficiently immunoprecipitates both DPF2 isoforms, ChIP-seq peaks were detected across the mouse genome. Peaks scoring >15 by HOMER were collected for further analysis. Comparing the data from cells expressing only one or the other isoform, we searched for differential ChIP peaks that would indicate preferential binding by either DPF2-S or DPF2-L. At sites that exhibit equal occupancy by the two DPF2 isoforms, other factors presumably determine BAF recruitment.

Interestingly, we found 445 genomic loci preferentially bound by DPF2-S, and 721 genomic loci preferentially bound by DPF2-L in mouse ESCs **(Figure 5A)**. We overlapped these differentially bound DPF2 peaks with enhancer elements identified as associated with a constellation of transcription factors in ESCs (Whyte et al., 2013). We found that 51 (∼12%) DPF2-S preferential sites and 69 (∼10%) DPF2-L preferential sites overlapped with predicted enhancer elements **(Figure 5B)**. We also overlapped the differential DPF2 ChIP peaks with ChIP peaks of the core BAF complex subunit BRG1 in mouse ESCs. This identified 168 (∼38%) DPF2-S preferential sites and 233 (∼33%) DPF2-L preferential sites that overlap with BRG1 binding sites, indicating that both DPF2-S and DPF2-L are binding as part of the BAF complex **(Figure 5C)**.

**Figure 5.**
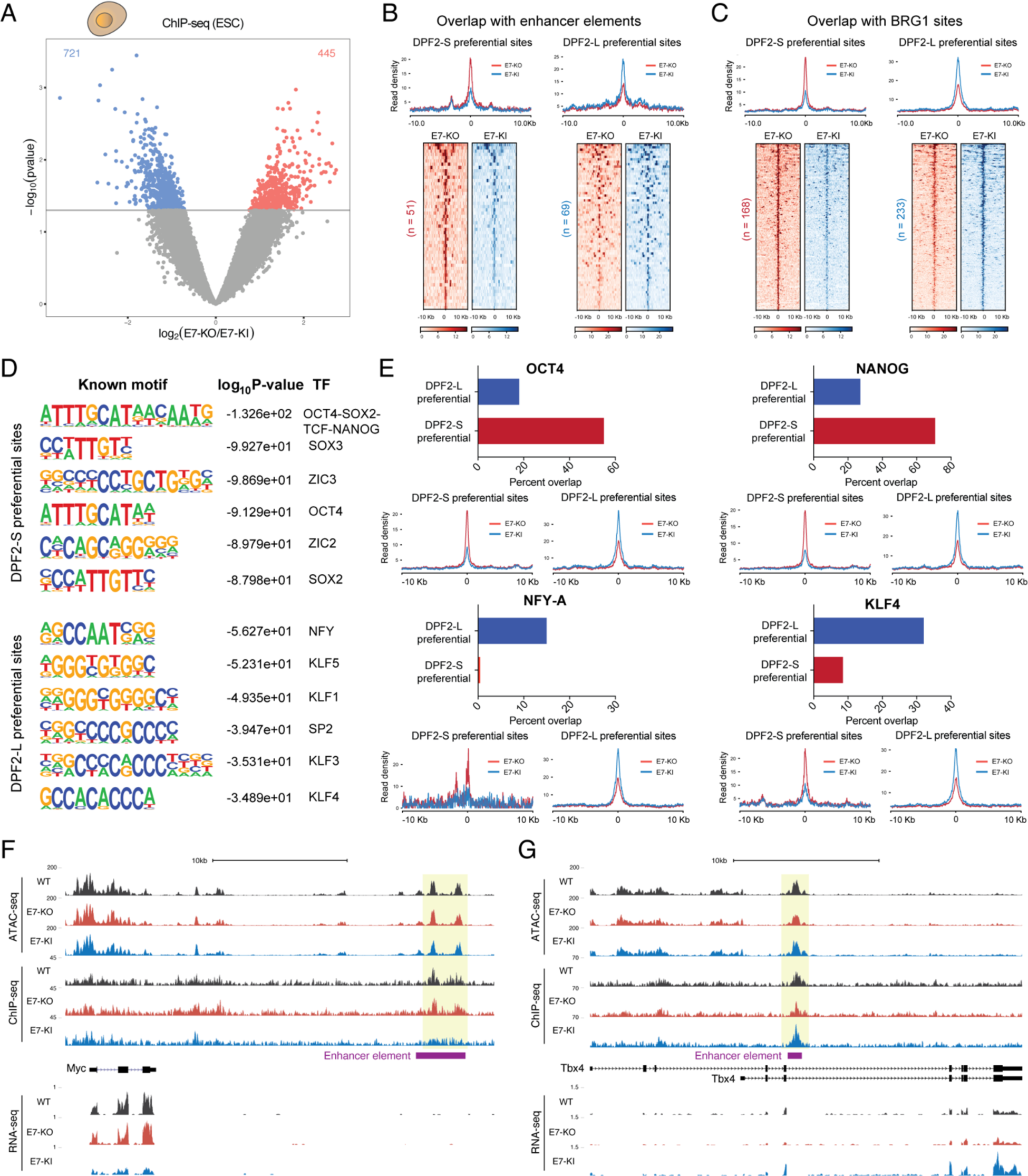
DPF2-S preferentially binds to genomic regions associated with stem cell pluripotency transcription factors in ESCs. **(A)** Volcano plot showing differential chromatin binding of DPF2-S and DPF2-L detected by ChIP-seq in mouse ESCs. Significant differential binding was filtered by 1.5-fold changes (*P*-value < 0.05) between E7-KO (expressing DPF2-S) and E7-KI (expressing DPF2-L). DPF2-S and DPF2-L preferential binding sites are labeled in red and blue dots, respectively. **(B-C)** Upper panel shows metaplot of average ChIP-seq signal intensities for DPF2-S and DPF2-L preferential binding sites centered upon the enhancer elements. (B) and known BRG1 binding sites (C) in ESCs. Lower panel shows normalized tag density profiles of preferential binding sites that overlap with predicted enhancers and BRG1 sites. Total number of preferential binding sites that overlap is indicated. **(D)** Enrichment of sequence motifs determined by HOMER analysis of the DPF2-S and DPF2-L preferential binding sites in ESCs. Log10 *P-*values and corresponding transcription factors are listed. **(E)** Percent overlap of the DPF2-S and DPF2-L preferential binding sites with known OCT4, NANOG, NFY-A, and KLF4 binding sites as determined by ChIP in ESCs. Bottom panel shows metaplots of average signal intensities for DPF2-S and DPF2-L preferential binding sites that overlap with the indicated transcription factor binding sites. **(F-G)** Genome browser tracks of ATAC-seq, ChIP-seq, and RNA-seq data at the *Myc* and *Tbx4* loci in wild-type (WT), DPF2-S expressing (E7-KO), and DPF2-L expressing (E7-KI) ESCs. Annotated ESC enhancer elements with differential DPF2-S and DPF2-L binding are highlighted in yellow. See also **Figures S4, S5**.

To confirm that both DPF2 isoforms were bound within full BAF complexes, we engineered ESC lines expressing either Flag-tagged DPF2-S or DPF2-L. We then immunoprecipitated each protein from either the soluble nucleoplasmic fraction (NP) or the high molecular weight (HMW) chromatin fraction of isolated nuclei and characterized the isolated proteins by SDS PAGE (Damianov et al., 2016) **(Figure S4A)**. As expected, the majority of the BAF complexes were found in the chromatin fraction. The major BAF subunits were all coimmunoprecipitated at similar stoichiometry to DPF2 indicating that the majority of DPF2 is associated with BAF **(Figures S4B-C)**. We did not observe major differences in the composition of BAF complexes pulled down by DPF2-S or DPF2-L, suggesting that the two isoforms assemble into similar complexes in ESCs.

We searched for known motifs enriched within the preferential binding sites for the DPF2-S and -L isoforms using the HOMER motif analysis algorithm (Heinz et al., 2010). Sequences enriched in the DPF2-S preferential binding sites were rich in A & T nucleotides and included motifs predicted to bind the pluripotency transcription factors OCT4, SOX2, and NANOG, as well as SOX3, ZIC2, and ZIC3 **(Figure 5D)**. Interestingly, both ZIC2 and ZIC3 were upregulated in DPF2-S expressing ESCs and have been previously shown to be required for maintenance of pluripotency, and ZIC3 also plays a role in establishing left-right asymmetry in the developing embryo (Herman and El-Hodiri, 2002; Lim et al., 2007; Luo et al., 2015). In contrast to DPF2-S, the enriched motifs within the DPF2-L preferential binding sites were high in G & C nucleotides and were predicted to bind a different set of TFs, including NFY, KLF1, KLF3, KLF4, KLF5, and SP2 **(Figure 5D)**. In addition to known binding motifs, we also identified *de novo* motifs enriched in loci preferentially bound by each of the two DPF2 isoforms **(Figure S5A)**. Some of these were again predicted binding sites for the pluripotency transcription factors or other proteins, and some did not correspond to known binding motifs.

To test whether the observed motif enrichments indicated transcription factor binding, we overlapped the preferential binding sites for DPF2-S and -L with known binding sites of the pluripotency TFs OCT4, NANOG, SOX2, and KLF4 (Chronis et al., 2017), as well as with NFY-A, identified in earlier ChIP-seq studies of mouse ESCs (Oldfield et al., 2014). Strikingly, ∼55%, ∼71%, and ∼51% of the DPF2-S preferential binding sites overlapped with known OCT4, NANOG, and SOX2 binding sites respectively, while the overlap with of the DPF2-L preferential binding sites was much lower (∼18%, ∼27%, and ∼23%) **(Figures 5E, S5B, S5D)**. Conversely, <1% and ∼9% of the DPF2-S preferential binding sites overlapped with mapped NFY-A and KLF4 binding sites respectively, while ∼15% and ∼32% of the DPF2-L preferential binding sites overlapped with sites bound by these factors, respectively **(Figures 5E, S5C)**. Binding sites for the paralogs NFY-B and NFY-C also overlapped with DPF2-L preferential sites (∼8% for both) but not with DPF2-S sites **(Figures S5G-H)**. Interestingly, HDAC1 and c-MYC binding sites showed substantial overlap with DPF2-L preferential binding sites (∼68% and ∼17%, respectively) compared to DPF2-S preferential sites (∼28% and <1%, respectively) **(Figures S5E-F)**. Taken together, these data indicate that BAF complexes containing DPF2-S and DPF2-L are recruited to distinct genomic loci occupied by different sets of transcription factors in ESCs.

To examine chromatin accessibility across the genome in DPF2-S or DPF2-L expressing ESCs, we performed transposase-accessible chromatin with sequencing assays (ATAC-Seq). This identified 477 genomic regions to be more accessible in DPF2-S expressing ESCs and 589 regions that were more open in DPF2-L expressing cells. These differentially accessible chromatin sites largely did not overlap with the DPF2-S or -L preferential binding sites identified by ChIP-seq, and their enriched motifs were different from those in the ChIP-seq data **(Figures S7A-B)**. The relatively small differences in transposase accessibility between cells expressing the two DPF2 isoforms indicates that they are not causing large scale changes in open versus closed chromatin. Instead, their interaction with different transcription factors may induce local changes in chromatin accessibility.

The alternative DPF2 isoforms were both seen to have effects on the expression of individual genes. For example, expression of the oncogene *Myc* was reduced in E7-KI ESCs (expressing DPF2-L), relative to both E7-KO and WT ESCs (expressing DPF2-S). We identified a preferential binding site for DPF2-S located ∼22 kb downstream of the *Myc* gene within an identified enhancer element active in mouse ESCs **(Figure 5F)**. ATAC-seq data did not indicate differences in the chromatin accessibility of this region. Instead, the increased binding of DPF2-S in this locus correlates with a positive effect on the expression of *Myc*. Similarly, *Tbx4* (T-Box Transcription Factor 4) gene is upregulated in E7-KI ESCs compared to E7-KO and WT ESCs. A known ESC enhancer element within a *Tbx4* intron shows higher binding of DPF2-L than DPF2-S, consistent with DPF2-L stimulating expression of Tbx4 **(Figure 5G)**. Overall, the data indicate that BAF complexes containing DPF2-S and -L differentially bind to specific genomic loci to regulate gene expression in stem cells, including genes associated with maintaining stem cell pluripotency.

### DPF2-S and -L differentially target new regulatory regions in NPCs

To examine the targeting of the two DPF2 isoforms in neural progenitor cells, we performed ChIP-seq in NPC’s 21 days after their derivation from embryoid bodies **(Figure 3D)**. Performing similar analyses to the ESC, we identified 491 genomic loci where DPF2-S preferentially binds over DPF2-L in NPC, and 385 genomic loci showing preferential binding for DPF2-L **(Figure 6A)**. We again overlapped these differential chromatin binding sites with previously described enhancer elements in NPCs (Creyghton et al., 2010). In contrast to the ESC enhancers that were defined by the binding of particular transcription factors, these NPC enhancers were predicted by the presence of H3K4 monomethyl and H3K27 acetyl modifications indicating loci of potentially active enhancers in these cells. Interestingly, 140 (∼29%) of the DPF2-S preferential peaks localized within NPC enhancer elements. However, only 3 (<1%) of the DPF2-L preferential binding sites mapped to predicted enhancers, indicating that DPF2-S is different from DPF2-L in its targeting to NPC enhancers **(Figure 6B)**.

**Figure 6.**
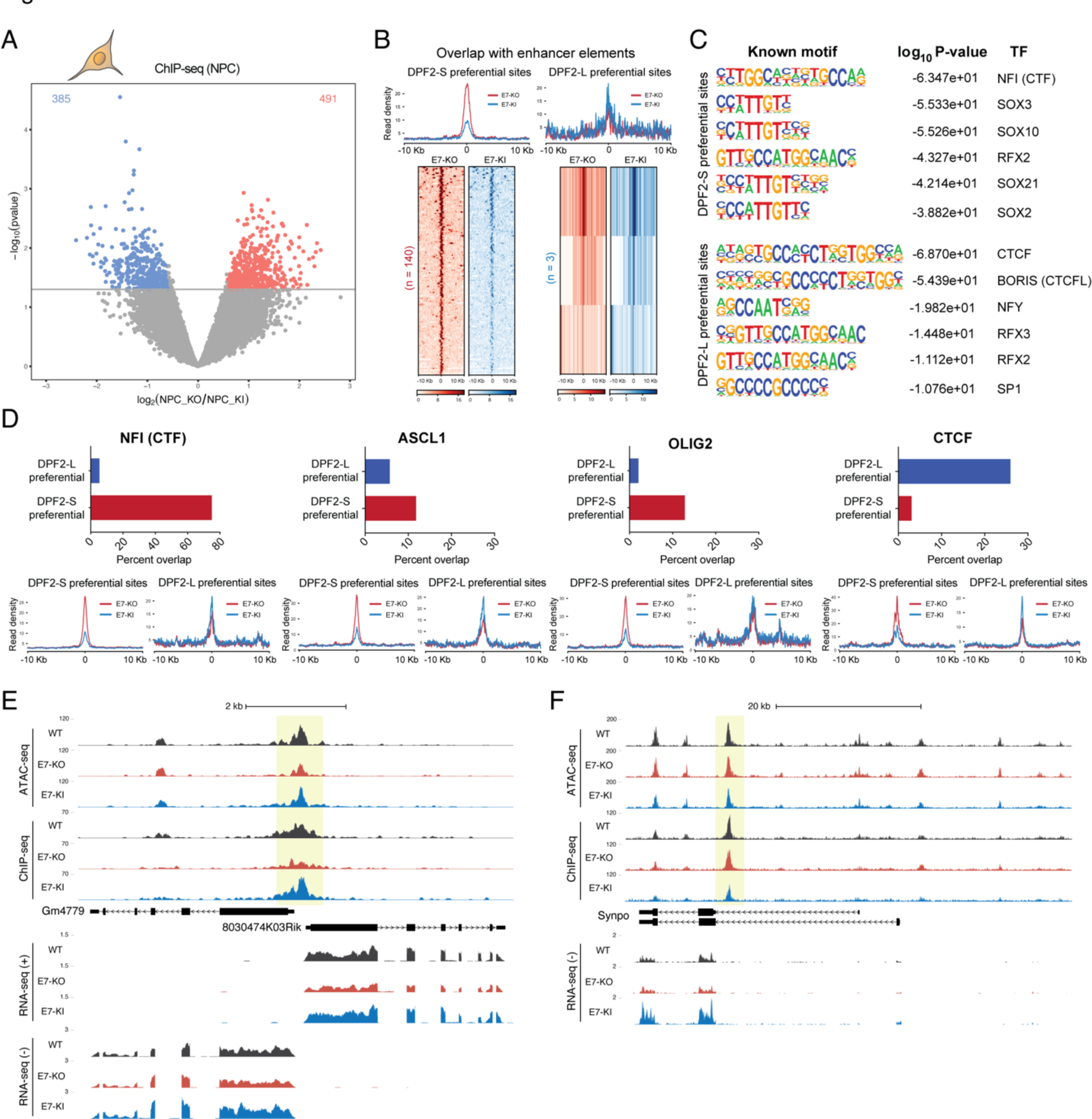
DPF2-S and -L preferentially bind to genomic regions associated with NFI or CTCF transcription factors in NPCs. **(A)** Volcano plot showing differential chromatin binding by DPF2-S and DPF2-L in mouse ESC-derived NPCs. Significant differential binding was cutoff at a 1.5-fold change (*P*-value < 0.05) between E7-KO and E7-KI cells. DPF2-S and DPF2-L preferential binding sites are labeled in red and blue dots, respectively. **(B)** Upper panel shows metaplot of average signal intensities for DPF2-S and DPF2-L preferential binding sites overlapping with predicted enhancer elements in NPCs. Lower panel shows normalized tag density profiles of preferential binding sites overlapping with predicted enhancers. **(C)** Enriched motifs identified by HOMER in DPF2-S and DPF2-L preferential binding sites in NPCs. Corresponding Log10 *P-*values and transcription factors are listed. **(D)** Percent overlap of DPF2-S and DPF2-L preferential binding sites with known NF1, ASCL1, OLIG2, and CTCF binding sites identified by ChIP-seq in NPCs. Bottom panels show metaplots of average signal intensities for DPF2-S and DPF2-L preferential binding sites overlapping with the transcription factor binding sites indicated above. **(E-F)** Genome browser tracks of ATAC-seq, ChIP-seq, and RNA-seq data at the *Gm4779*, *8030474K03Rik* (E), and *Synpo* (F) loci in wild-type (WT), DPF2-S expressing (E7-KO), and DPF2-L expressing (E7-KI) NPCs. Regions with differential DPF2-S and DPF2-L binding are highlighted in yellow. See also **Figure S6**.

Assessing the enriched sequence motifs in the two groups of differential binding sites also identified differences in the targeting of the two isoforms. The top DPF2-S motifs included transcription factor binding sites such as NFI, SOX3, SOX10, RFX2, SOX21, and SOX2. In contrast, the DPF2-L preferential motifs match binding motifs for CTCF, BORIS (CTCFL), NFY, RFX2, and SP1 **(Figure 6C)**. We also identified *de novo* motifs enriched in the loci preferentially bound by DPF2-S or -L **(Figure S6A)**. Similar to the known motif analysis, the top *de novo* motif for DPF2-S preferential sites was an NFI binding site, whereas the top *de novo* motif for DPF2-L is predicted to bind the CTCF paralog BORIS (CTCFL).

Examining ChIP-seq data for some of these factors in NPC (Beagan et al., 2017; Mateo et al., 2015; Nishi et al., 2015; Wapinski et al., 2013), we found that ∼75% of the DPF2-S preferential sites overlapped with known NFI binding sites, while only ∼6% of the DPF2-L sites overlapped with NFI **(Figures 6D, S6B)**. Conversely, only ∼3% of the DPF2-S preferential sites overlapped with CTCF binding sites, while ∼26% of the DPF2-L preferential sites showed overlap with CTCF sites **(Figures 6D, S6E)**. Interestingly, the DPF2-S and DPF2-L preferential sites showed more limited overlap with the binding sites of the NPC specific transcription factors ASCL1 and OLIG2, with DPF2-S exhibiting higher overlap than DPF2-L **(Figures 6D, S6C-D)**.

ATAC-seq analyses in NPCs elicited similar results to ESCs. We identified 472 genomic regions to be more accessible in DPF2-S-expressing NPCs, and 315 regions to be more accessible in DPF2-L-expressing NPCs. Again, these regions largely did not overlap with the DPF2-S or -L preferential binding sites identified by ChIP-seq **(Figures S7C-D)**.

Depending on the target gene, the binding of the DPF2 isoforms may regulate expression in opposing directions. For example, the Synaptopodin gene (*Synpo*) is upregulated in the E7-KI NPCs compared to both E7-KO and WT NPCs **(Figure 6F)**. ChIP-seq revealed a locus within the Synpo gene where DPF2-S binds more strongly than DPF2-L, suggesting either downregulation of the gene by DPF2-S or a greater stimulatory activity by the limited amount of DPF2-L. Also upregulated in E7-KI NPCs compared to the E7-KO are two uncharacterized divergent genes Gm4779 and 8030474K03Rik **(Figure 6E)**. In this case, the DPF2-L isoform preferentially binds with the common promoter region shared by these genes, consistent with DPF2-L stimulating their expression. ATAC-seq revealed that chromatin accessibility in this region is lower in the E7-KO NPCs than the E7-KI. We also observed intermediate levels of expression of these genes in the WT NPCs with intermediate levels of DPF2 binding, perhaps reflecting the expression of both isoforms in these cells (∼80% DPF2-S and ∼20% DPF2-L).

Overall, these data indicate that in neuronal progenitor cells DPF2-S and -L are targeted to different genomic loci that are associated with the proteins NFI or CTCF respectively.

### DPF2-S binds to chromatin sites with enhancer specific modifications, and DPF2-L binds to sites enriched for promoter modifications

To further analyze the sites of differential DPF2-S and -L binding, we intersected their binding sites with previously annotated ESC chromatin states defined by ChromHMM (Chronis et al., 2017; Ernst and Kellis, 2017). ChromHMM chromatin states correspond to the combinatorial presence or absence of multiple histone modifications within their chromosomal context. The states are associated with different classes of genomic elements such as enhancers of varying activity levels, active and poised promoters, transcribed and repressed regions, and regions with minimal or no signal for any histone mark. To confirm the associations identified by ChromHMM, we measured the levels of diagnostic histone modifications around the binding sites of DPF2-S and -L. ChromHMM indicated that in ESCs, DPF2-S sites show strong overlap with regions having the characteristics of active, weak, and poised enhancers (**Figure 7A**).

**Figure 7.**
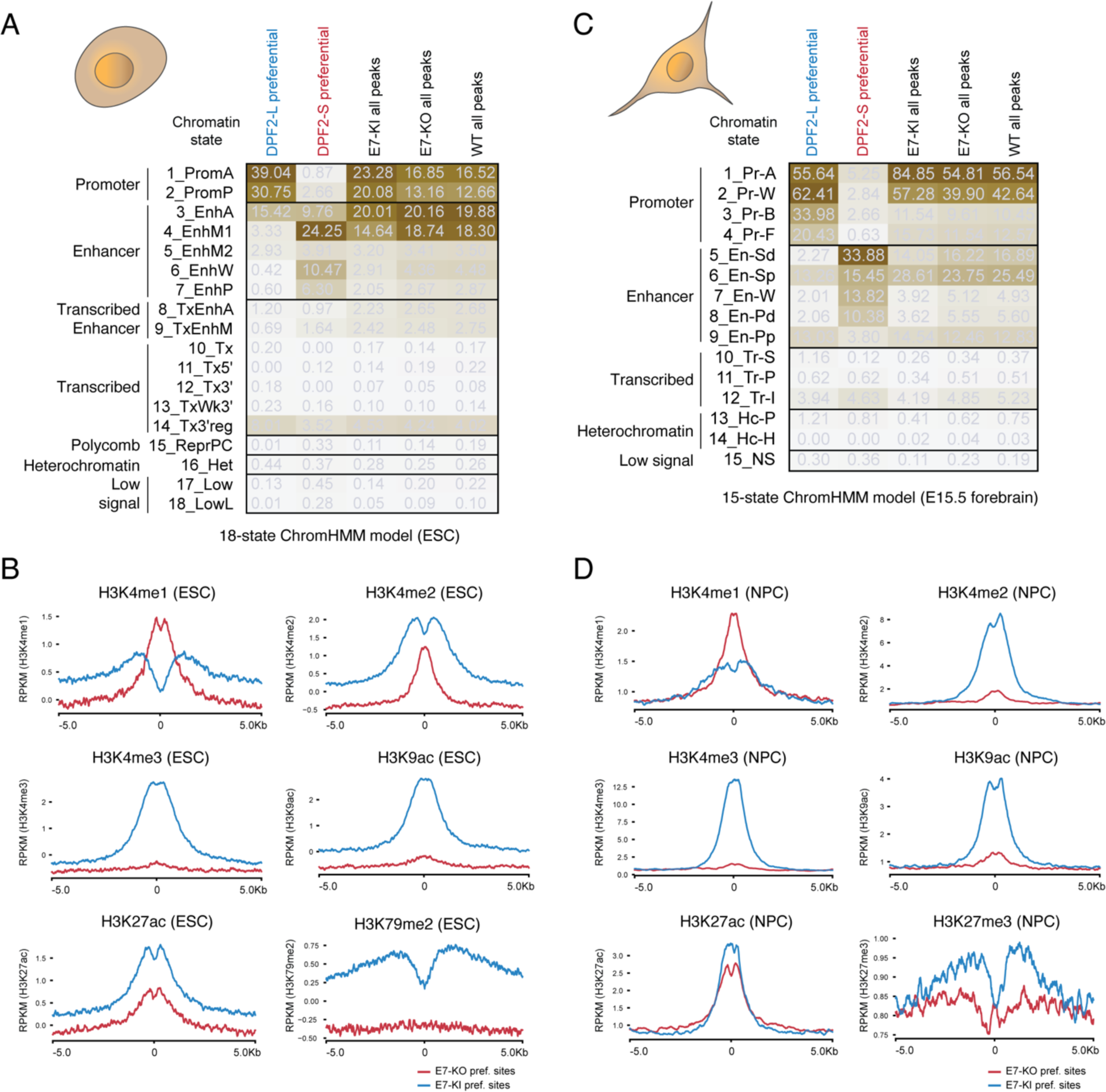
DPF2-S binds to chromatin sites with enhancer specific states, and DPF2-L binds to sites enriched for promoter states in both ESCs and NPCs. **(A)** Analysis with an 18-state ChromHMM model (Chronis et al., 2017) of DPF2-S and DPF2-L preferential binding sites in ESCs. The rows represent different chromatin states and are labeled by their representative mnemonics. Columns show the fold enrichments (ChromHMM emission probabilities) of these states for each set of DPF2 ChIP sites, including DPF2-S and DPF2-L preferential binding sites, colored from highest to lowest. **(B)** Metaplots of signal intensities for H3K4me1, H3K4me2, H3K4me3, H3K9ac, H3K27ac, H3K79me2 chromatin modifications measured by ChIP-seq in ESCs near the DPF2-S and DPF2-L preferential binding sites. **(C)** Analysis with a mouse E15.5 forebrain-specific 15-state ChromHMM model (Gorkin et al., 2020) of DPF2-S and DPF2-L preferential binding sites in NPCs. **(D)** Metaplots of signal intensities for H3K4me1, H3K4me2, H3K4me3, H3K9ac, H3K27ac, H3K27me3 chromatin modifications in NPCs near the DPF2-S and DPF2-L preferential binding sites. See also **Figure S7**.

Notably, the enhancer mark H3K4me1 was highly enriched at DPF2-S sites but depleted at DPF2-L preferential sites **(Figures 7B)**. In contrast, DPF2-L sites show chromatin characteristics of promoter regions. These sites showed high levels of the histone mark H3K4me3 (a promoter mark), as well as the marks H3K9ac (found in active regions—promoter and enhancer), H3K27ac (enhancers), and H3K79me2 (transcribed regions) **(Figures 7A-B)**. The H3K4me2 mark was found at both DPF2-S and -L preferential sites, but was distributed differently. At DPF2-S sites, this H3K4me2 modification showed a single enrichment peak centered upon the ChIP-seq peak, whereas for DPF2-L sites H3K4me2 was most strongly enriched in the flanking regions upstream and downstream of the ChIP-seq peak **(Figure 7B)**.

We next analyzed the DPF2 ChIP-seq data in ESCs using a recently developed pan-cell type (‘universal’) chromatin state annotation of the mouse genome trained on 901 epigenomic datasets profiling histone modification and open chromatin signals across many cell and tissue types (denoted the full-stack annotation (Vu and Ernst, 2022)). Consistent with the results from the 18-state ChromHMM model, the full-stack annotation also showed DPF2-S preferentially binding to enhancer elements with the chromatin states mEnhA7-11 **(Figure S7E)**. These states are characterized by high levels of enhancer-associated marks (H3K4me1, H3K27ac, and H3K9ac). The DPF2-S binding sites are also enriched for the ESC-specific weak enhancer state mEnhWk1 state (See Supplemental Table S6 for a list of chromatin marks that define each state in the stacked ChromHMM model). The stacked model again identified DPF2-L binding sites as associated with promoter and transcription start site (TSS)-associated chromatin states, with the strongest enrichment for the constitutive active TSS states (mTSS1-3) **(Figure S7E)**. When applied to the ESC ChIP data, both ChromHMM annotations associated DPF2-S binding with enhancers, and DPF2-L binding with promoters.

To examine the chromatin landscape of DPF2 binding in NPCs, we used a 15-state ChromHMM model that captures combinatorial patterns of histone modifications across a range of mouse tissues and developmental stages (Gorkin et al., 2020). We used a forebrain-specific annotations at developmental day E15.5, a stage where most cells are in a neural progenitor like state. In this model, DPF2-S preferential binding sites from NPCs were enriched for multiple enhancer states, including En-Sd (Enhancer, Strong TSS-distal), En-W (Enhancer, Weak TSS-distal), and En-Pd (Enhancer, Poised TSS-distal). Another enhancer state En-Sp (Enhancer, strong TSS-proximal) was enriched in both the DPF2-S and DPF2-L preferential sites, while the En-Pp state (Enhancer, Poised TSS-proximal) was enriched in DPF2-L preferential sites **(Figure 7C)**. Confirming some of the individual marks that define these chromatin states, we found that DPF2-S sites were distinguished by high levels of H3K4me1 and H3K27ac, characteristic of active enhancers in NPCs **(Figure 7D)**. This NPC ChromHMM model also associated DPF2-L preferential sites with four promoter states Pr-A (Promoter, Active), Pr-W (Promoter, Weak/Inactive), Pr-B (Promoter, Bivalent), and Pr-F (Promoter, Flanking) **(Figure 7C)**. These sites show strong enrichment for the histone marks H3K4me2, H3K4me3, and H3K9ac compared to DPF2-S binding sites **(Figures 7D)**.

The stacked ChromHMM model (Vu and Ernst, 2022) gave similar results for the NPC ChIP-seq data. DPF2-S bound to sites enriched for the chromatin enhancer states mEnhA1-4 and mEnhA9-A11, representing active and weak to moderate enhancers respectively. DPF2-L sites were again most strongly enriched for promoter and transcription start site states including the constitutive active TSS states mTSS1-3 **(Figure S7F)**. Consistent with our finding that DPF2-L sites often contain CTCF binding sites in NPCs **(Figures 6D)**, DPF2-L sites were enriched for the mOpenC6 and mOpenC7 states that represent candidate insulators with high CTCF binding (Vu and Ernst, 2022). In sum, the chromatin state analyses in both cell types identify DPF2-S targeting to enhancer regions, and DPF2-L targeting to promoters and transcriptional start sites, as well as to insulator elements bound by CTCF.

Altogether the data show that the change in DPF2 spliced isoforms redirects chromatin remodeling BAF complexes to different genomic loci serving different genomic functions.

## DISCUSSION

### The PTBP1 regulatory switch creates a new neuronal chromatin remodeling factor

We found that during neuronal development, splicing of the BAF complex subunit DPF2 is switched from its canonical DPF2-S isoform to a neuronal DPF2-L protein isoform. The *Dpf2*-L mRNA contains a new exon 7 that modifies its C2H2 zinc-finger domain. We show that the splicing regulator PTBP1 represses exon 7 splicing through binding to CU-rich elements in its flanking introns. This allows Exon 7 to be induced early in neuronal differentiation, when PTBP1 is depleted **(Figure 8)**.

**Figure 8:**
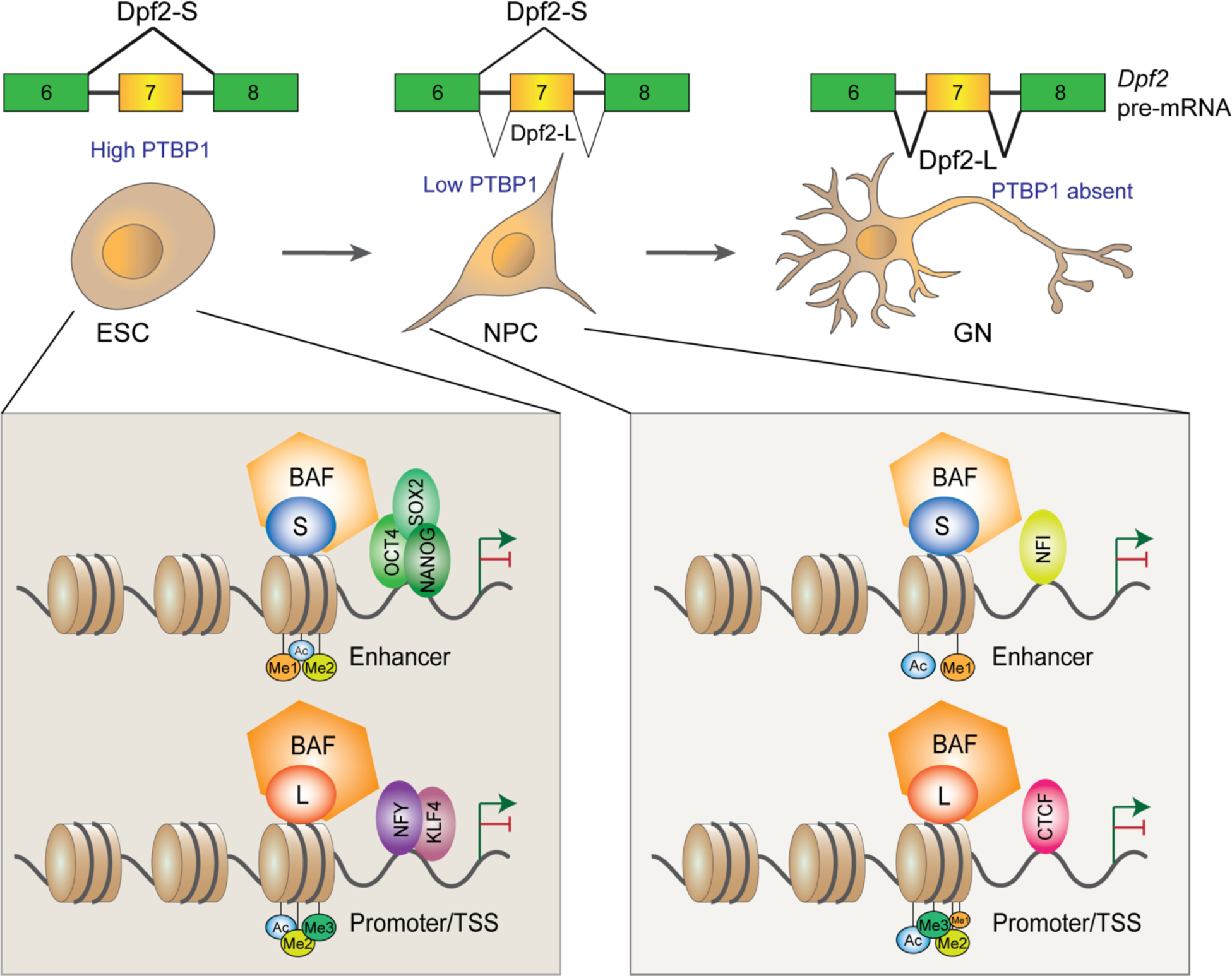
Functional changes in *Dpf2* spliced isoforms during ESC maintenance and neuronal differentiation. *Dpf2* exon 7 is strongly repressed by PTBP1 in ESCs leading to DPF2-S expression. Depletion of PTBP1 early in neuronal differentiation induces exon 7 inclusion and switches expression to DPF2-L isoform. In ESCs, DPF2-S directs BAF to enhancers bound by pluripotency transcription factors including OCT4, NANOG, and SOX2. Forced expression of DPF2-L in ESC redirects BAF to promoter/TSS specific regions bound by NFY and KLF4. In NPCs, DPF2-S targets BAF to enhancer loci bound by the NFI transcription factor. In contrast, DPF2-L in NPC recruits BAF to promoters and TSS bound by the genomic insulator transcription factor CTCF. Distinct histone modifications define these enhancers or promoter/TSS-specific chromatin states and are enriched at the specific sites of DPF2-S and DPF2-L binding. Thus, splicing regulatory events during development elicit wide ranging impacts on chromatin organization and transcription.

Prior studies have shown that PTBP1 maintains a program of nonneuronal splicing in ESCs and neuronal progenitor cells (Keppetipola et al., 2012; C. K. Vuong et al., 2016). When neuronal progenitors exit mitosis and begin to differentiate into neurons, PTBP1 expression is repressed through the action of the microRNA miR124 (Makeyev et al., 2007). The transition to a neuronal phenotype is reinforced by regulatory loops where the loss of PTBP1 relieves blocks both to miR124 biogenesis and to miR124-mediated repression of the transcriptional regulators, SCP1 and CoREST (Cheng et al., 2009; Makeyev et al., 2007; Visvanathan et al., 2007; Yeom et al., 2018). The loss of PTBP1 also causes the induction of its paralog PTBP2 due to the loss of repression of a *Ptbp2* exon, whose skipping leads to nonsense mediated mRNA decay (Boutz et al., 2007).

The two PTB proteins have extensive impacts on the posttranscriptional regulatory programs of development (Boutz et al., 2007; Keppetipola et al., 2012; C. K. Vuong et al., 2016; J. K. Vuong et al., 2016). Through binding to CU-rich elements in pre-mRNAs, these proteins repress splicing of many exons, but can also stimulate splicing of certain exons and cause retention of some introns (Hamid and Makeyev, 2017; Llorian et al., 2010; Yap et al., 2012; Yeom et al., 2021). Some exons are regulated by both proteins and maintain their repression through the early switch from PTBP1 to PTBP2 (Li et al., 2014; Licatalosi et al., 2012; Zheng et al., 2012). Other exons that are more sensitive to PTBP1 than PTBP2 shift their splicing earlier in development when the expression of the two regulators is switched (Boutz et al., 2007; Linares et al., 2015). This differential sensitivity may be due in part to these exons requiring dimerization of the PTBP1 protein to be repressed, and the lack of dimerization seen for PTBP2 (Ye et al., 2023). Isoforms repressed by PTBP1 early in development affect functions such as axonogenesis, cell polarity, reduced apoptotic potential, and transcriptional programs of early neurons (Lin et al., 2020; Linares et al., 2015; M. Zhang et al., 2019).

Recent studies reported that depletion of PTBP1 or co-depletion of PTBP1 and PTBP2 was sufficient to induce the transdifferentiation of cells such as fibroblasts or astrocytes into fully mature neurons (Maimon et al., 2021; Qian et al., 2020; Xue et al., 2013; Zhou et al., 2020). This has been employed in strategies to restore neuronal cell numbers in models of neurodegeneration, although other groups have not replicated these findings (Chen et al., 2022; Hoang et al., 2022; Wang et al., 2021). In mouse ESC, we do not see full neuronal differentiation after PTBP1 and/or PTBP2 depletion, although the cells now express multiple neuronal markers (Linares et al., 2015; Yeom et al., 2018). These different observations in ESC may arise from the programs maintaining pluripotency in ESC adding controls on PTBP1 targets that are absent in the fibroblasts used in the transdifferentiation studies.

Exon 7 of *Dpf2* is induced early in neuronal development coincident with the switch from PTBP1 to PTBP2. The change in DPF2 proteins extends the known impact the PTBP switch to chromatin regulation within the differentiating neurons. Interestingly, other BAF complex subunits are substituted with neuronal paralogs at a similar time in development (Kadoch and Crabtree, 2015; Lessard et al., 2007; Narayanan and Tuoc, 2014; Ronan et al., 2013; Staahl et al., 2013). These neuronal BAF subunits alter functions such as cell cycle exit in the differentiating cells (Braun et al., 2021). Mutation or misregulation of BAF subunits are also linked to a number of neurodevelopmental disorders (Ronan et al., 2013; Sokpor et al., 2017). Human mutations in DPF2 itself are associated with Coffin Siris Syndrome whose pleiotropic phenotypes include intellectual disabilities and neurological dysfunction. These defects associated with Dpf2 mutation and the high conservation of *Dpf2* exon 7 across vertebrate species indicate an important function in the brain. It will be interesting to explore the neuronal function of exon 7 in genome-edited mice.

### The switch in *Dpf2* splicing retargets BAF complexes from enhancers to promoters

We observed that expression of DPF2-L in ESCs and neural progenitor cells (NPCs), or of DPF2-S in glutamatergic neurons (GNs) caused substantial shifts in gene expression from cells expressing the other isoform (**Figures 4, S3**). We further found that DPF2-S and DPF2-L preferentially bound chromatin sites in ESCs and NPCs characterized by distinct histone marks and transcription factor binding. In ESCs, DPF2-S exhibited enhanced binding to chromatin sites targeted by core pluripotency transcription factors including OCT4, SOX2, and NANOG **(Figures 5E, S5B, S5D)**. This is consistent with studies showing that DPF2 binding sites in ESCs overlap with OCT4 and SOX2 binding sites (W. Zhang et al., 2019), as well as with reports that DPF2, along with other BAF complex subunits are components of an OCT4 protein network required for ESC self-renewal (Pardo et al., 2010; van den Berg et al., 2010). Notably, DPF2-L was targeted differently in ESC, and its preferential binding sites exhibited limited overlap with the core pluripotency transcription factors. Instead, the DPF2-L sites were enriched for motifs of factors such as NFY and KLF4 **(Figures 5E, S5C)**. Switching from DPF2-S to DPF2-L in ESCs did not affect the levels of OCT4, SOX2, and NANOG transcripts, but expression of the transcription factors *Zic2*, *Zic3*, and *Myc* was reduced by DPF2-L. MYC can potentiate the action of the pluripotency transcription factors in ESCs (Fagnocchi et al., 2016; Rahl et al., 2010). ZIC2 and ZIC3 are C2H2-type zinc finger proteins required for the maintenance of ESC pluripotency, and are directly regulated by core pluripotency TFs OCT4, SOX2, and NANOG (Lim et al., 2007; Luo et al., 2015). ZIC3 also plays a role in establishing a proper left-right axis and in midline neural patterning during embryonic development (Herman and El-Hodiri, 2002). In Xenopus, ZIC3 functions as a left-side determinant after laterality is established in early embryos (Kitaguchi et al., 2000). ZIC3 controls expression of *xnr1* (nodal-related 1) and *pitx2*, which are key determinants of organ asymmetry (Kitaguchi et al., 2002). Intriguingly, we observed increased expression of *Nodal* and *Pitx2*, as well as the left-right determination factors *Lefty1* and *Lefty2*, in DPF2-S expressing ESCs. These expression changes in asymmetry genes may result from the higher MYC, ZIC3 and/or ZIC2 levels in these cells **(Figures 4A-B)**. It will be interesting to explore whether DPF2 plays a role in asymmetry generation in the early embryo.

In neuronal progenitor cells the two DPF2 isoforms were targeted differently than in ESCs. The most enriched motif within the DPF2-S preferential sites in these cells was the binding site for the nuclear factor 1 (NFI, also called NFI/CTF). NFI/CTF is member of the CAAT box binding family of transcription factors that control developmentally regulated genes. NFI also acts at intra-chromosomal boundary elements affecting both transcription and DNA replication (Gaussin et al., 2012). In mice, the loss of NFI paralogs NFIa, NFIb, NFIx leads to multiple abnormalities of brain development (das Neves et al., 1999; Driller et al., 2007; Shu et al., 2003). In contrast, the most enriched motif in the DPF2-L preferential sites of NPCs was a binding site for the CCCTC binding factor (CTCF). CTCF is a zinc finger protein that regulates enhancer-promoter interactions by chromatin looping, acts as an insulator protein that blocks enhancers from activating promoters, and forms ‘CTCF loop anchors’ at the base of transcriptionally active chromatin loops (Arzate-Mejía et al., 2018; Fudenberg et al., 2016; Recillas-Targa et al., 2002; Sanborn et al., 2015). CTCF restricts premature neurogenesis by affecting the cell survival, proliferation, and controlled differentiation of neuronal progenitors in early embryonic development. Depletion of CTCF in the NPCs of E11 mouse brain caused apoptosis and lethality at birth (Watson et al., 2014). In binding to sites occupied by CTCF in NPCs **(Figures 6D, S6E)**, DPF2-L may alter expression of multiple genes within a chromatin loop and thereby affect many aspects of neuronal differentiation. It will be interesting to examine whether DPF2-L binding sites in mature brain also coincide with CTCF binding.

The preferential binding sites of DPF2-S and DPF2-L were also distinguished by their chromatin environments, as defined by ChromHMM. We used an 18-state ChromHMM model for MEFs, pre-iPSCs, and ESCs (Chronis et al., 2017), a 15-state model for multiple tissues across mouse development (Gorkin et al., 2020), and a universal 100-state stacked model for all cell types (Vu and Ernst, 2022). These models all associated different chromatin states with the binding sites for the two DPF2 isoforms. In ESCs and NPCs, DPF2-S was seen to bind chromatin in enhancer specific states, while DPF2-L preferred binding near promoters and transcription start sites (TSS) **(Figures 7A, 7C)**. The different chromatin states were confirmed in ChIP-seq data where in ESCs we found enrichment of H3K4me1 and H3K4me2, and to a lesser extent H3K27ac at DPF2-S binding sites. This agrees with a report identifying DPF2 as a H3K4me1-associated protein at mammalian enhancers (Local et al., 2018). In contrast, DPF2-L binding sites in both ESCs and NPCs were enriched for H3K4me2, H3K4me3, H3K9ac, and H3K27ac but exhibited minimal presence of H3K4me1 **(Figures 7B, 7D)**. Thus, the induction of the Dpf2-L isoform allows the retargeting of BAF complexes to new chromatin sites as neurons mature.

### Significance and limitations of these studies

Our results connect two processes by which gene expression is altered during neuronal development: the induction of alternatively spliced isoforms by changes in the PTB proteins, and the exchange of BAF complex subunits to create neuron-specific chromatin modifiers. We identify a PTBP1-dependent BAF complex subunit that is specifically expressed in neurons and show that this alters the transcriptional profile in the cells where it is expressed. We find that this DPF2-L isoform can target the BAF complex to new genomic loci and to promoters rather than enhancers. It is not yet clear from these studies what direct molecular interactions are altered by the inclusion of exon 7. Further studies in mice will also be needed to assess the role of the DPF2-L regulatory program in brain development.

## ACKNOWLEDGEMENTS

We thank Hynek Wichterle and Tom Jessell for the HBG3 ES cell line, Reem Halabi and Marcus Tol for providing mouse RNA samples, and Wensheng Zhang for C-TAP-*Dpf2* clones. We thank James Wohlschlegel and Yasi Jami for mass spectrometry analysis. We thank Andrey Damianov and all the members of the Black Laboratory for their help and constructive discussions. We thank Kathrin Plath for valuable suggestions and discussions. This work was supported by NIH grant R35 GM136426 to DLB, a SEED grant from the Jonsson Comprehensive Cancer Center at UCLA to DLB, and an Innovation Award from the Broad Stem Cell Research Center at UCLA to DLB.

## AUTHOR CONTRIBUTIONS

M.N. and D.L.B. designed the research with advice from J.E., M.F.C., and S.T.S.; M.N. performed experiments and analyzed data with help from A.-C.F., K.-H.Y. and M.L.; C.-H.L., A.-C.F., A.E.D., H.V., and X.T. performed bioinformatic analyses; and M.N. and D.L.B. wrote the manuscript with input from the other authors.

## DECLARATION OF INTERESTS

D.L.B. has equity and serves on the board of directors for Panorama Medicine. This company did not contribute to or direct any of the research reported in this article. The authors declare no competing interests.

## MATERIALS AND METHODS

### Cell lines and tissue culture

Mouse HB9-GFP (HBG3) ESCs were gifts from H. Wichterle and T. Jessell at Columbia University. Cells were grown with ESC media consisting of DMEM (Fisher Scientific) supplemented with 15% ES-qualified fetal bovine serum (Life Technologies), ESC-qualified nucleosides (EMD Millipore), GlutaMAX (Life Technologies), non-essential amino acids (Life Technologies), 0.1 mM 2-Mercaptoethanol (Sigma-Aldrich), and 1000U/ml of ESGRO leukemia inhibitor factor (EMD Millipore) on 0.1% gelatin-coated dishes with CF1 mouse embryonic fibroblasts (Applied StemCell) as feeder cells.

### Construction of minigene reporters

We constructed mouse *Dpf2* minigene by inserting the genomic locus encompassing exon 6 to exon 8 in the pcDNA3.1+ vector using EcoRI and NotI sites. Artificial mutations were introduced in the minigene by Reverse Site-Directed Mutagenesis with Phusion High-Fidelity DNA Polymerase (NEB), followed by 5’ phosphorylation with T4 Polynucleotide Kinase (NEB) and self-ligation with T4 DNA Ligase (NEB).

### Knockdown of *Ptbp1* and *Ptbp2* in mouse ESCs

HBG3 ESCs were transfected with Silencer Select siRNAs against *Ptbp1* (Thermo Fisher Scientific; s72335, s72337) and *Ptbp2* (Thermo Fisher Scientific; s80148, s80149) using RNAiMax (Thermo Fisher Scientific) according to the manufacturer’s protocol. Silencer Negative Control siRNA #1 (Thermo Fisher Scientific) was used as negative control. ESCs were transfected with 20 nM of siRNAs followed by a repeated treatment after 24 hrs and harvested 72 hr post-transfection.

### CRISPR/Cas9-mediated genome editing in mouse ESCs

Guide RNAs (gRNAs) targeting *Dpf2* introns 6 and 7 were designed using the CRISPOR (http://crispor.tefor.net/) (Concordet and Haeussler, 2018) tool. For knocking out of *Dpf2* exon 7, RNP complex was assembled using 30 μM (pmol/μl) of Synthetic CRISPRevolution sgRNAs (Synthego) and 90 pmol Cas9 endonuclease 2NLS, *S. pyogenes* (Synthego) for 10 minutes at room temperature. Cell suspension containing ∼100k cells was electroporated using the Neon Transfection System (Fisher Scientific) with 1250V pulse voltage and 2 pulses with a pulse width of 20 ms. For knocking-in exon 7 cassette with modified 5’ and 3’ splice sites, we electroporated the same sgRNAs and Cas9 RNP complex with 20 pmol of a 761 nt ssDNA donor repair template (GenScript) harboring a 161 nt knock-in insert and 300 nt homology arms on both sides. After two days in culture, CRISPR-edited heterogenous pool of ESCs were genotyped for checking the presence of desired edits. Individual genome-edited clones were isolated by limiting dilution followed by clonal expansion. Cells with homozygous deletion of *Dpf2* exon 7 or homozygous knock-in of modified exon 7 cassette were genotyped by PCR. Cells with no deletion or insertion from the same heterogenous pool were used as controls.

### Differentiation of mouse ESCs to NPCs

HBG3 ESCs were differentiated into NPCs as previously described (Conti et al., 2005; Linares et al., 2015), with some modifications. Briefly, feeder-free ESCs were differentiated on 0.1% gelatin-coated dish in N2B27 media consisting of 1:1 mixture of DMEM/F12 (Fisher Scientific) and Neurobasal media (Life Technologies) supplemented with 0.5X B27 (without Vitamin A, Life Technologies), 0.5X N2 Supplement (Life Technologies), GlutaMAX (Life Technologies), 100 U/mL Penicillin-Streptomycin (Thermo Fisher Scientific), and 0.1 mM 2-Mercaptoethanol (Sigma-Aldrich). After growing in aggregate cultures for seven days, cells were trypsinized and single cells were passed through a 40 μm nylon strainer (Thermo Fisher Scientific). Single cells were grown in N2B27 media supplemented with 10 ng/ml of recombinant human EGF (PeproTech) and FGF-basic (PeproTech) for two days in non-gelatinized plates. Cells were then dissociated with TrypLE Express Enzyme (Thermo Fisher Scientific) and plated on poly-ornithine-(15 ug/ml, Sigma) and fibronectin-coated (1.5 ug/ml, Sigma-Aldrich) dishes in NPC media consisting of DMEM/F12 (Fisher Scientific) supplemented B27 (without Vitamin A, Life Technologies), GlutaMAX (Life Technologies), 10 ng/ml recombinant human EGF (PeproTech), 10 ng/ml recombinant human FGF-basic (PeproTech), and 0.1 mM 2-Mercaptoethanol (Sigma-Aldrich). Differentiated NPCs were maintained on poly-ornithine- and fibronectin-coated dishes in NPC media for several passages.

### Differentiation of mouse ESCs to GNs

HBG3 ESCs were differentiated to glutamatergic neurons (GNs) by a method adapted from previous studies (Gueroussov et al., 2015; Hubbard et al., 2013) with modifications. Briefly, feeder-free mouse ESCs were grown as aggregate culture for eight days (DIV-8) in differentiation media consisting of Glasgow’s MEM (GMEM) (Thermo Fisher Scientific) supplemented with 5% Knockout Serum Replacement (KSR) (Invitrogen), non-essential amino acids (Life Technologies), 1mM Sodium Pyruvate (Invitrogen), GlutaMAX (Life Technologies), and 0.1 mM 2-Mercaptoethanol (Sigma-Aldrich). Differentiating aggregates/embryoid bodies (EBs) were maintained at 37°C, 5% CO2 and 90% relative humidity with 50% media replaced by fresh media in every two days. After four days, the differentiation media was supplemented with 6 μM all-trans retinoic acid (RA, Sigma-Aldrich) until day 8. On day 8 (DIV 0), EBs/aggregates were dissociated with TrypLE Express Enzyme (Thermo Fisher Scientific) for 5 minutes at 37°C and dissociation was stopped by Trypsin inhibitor from Glycine max (Sigma-Aldrich). Single cells were then passed through a 40 μm nylon strainer (Thermo Fisher Scientific) and plated in N-2 media consisting of Neurobasal-A medium (Invitrogen) supplemented with N-2 Supplement (Life Technologies), GlutaMAX (Life Technologies), and 100 U/mL Penicillin-Streptomycin (Thermo Fisher Scientific) on Poly-D-lysine hydrobromide (Sigma-Aldrich) and Laminin (Sigma-Aldrich) coated dishes. Complete media changes were done 4 hours and 24 hours after plating. Two days later (DIV 2), N-2 media was replaced by B-27 media consisting of B-27 Serum-Free Supplement (Thermo Fisher Scientific), GlutaMAX (Life Technologies), and 100 U/mL Penicillin-Streptomycin (Thermo Fisher Scientific). Differentiating neurons underwent full media change in every 3 days intervals. On DIV-5, the media was supplemented with 10 μM 5’-fluoro-2’-deoxyuridine (Sigma-Aldrich) and 30 μM Uridine (Sigma-Aldrich) to select against any remaining glia until DIV-10.

### RNA extraction and RT-PCR

Total RNA was extracted from cell cultures using Trizol (Life Technologies) according to the manufacturer’s instructions. RNA was quantified using the Nanodrop-1000 spectrophotometer (Thermo Fisher) and 1 mg total RNA was used for each 20 ul reverse transcription reaction using SuperScript III RT enzyme (Life Technologies) followed by PCR reactions using the GoTaq Green Master Mix (Thermo Fisher Scientific). Bands were quantified using the ImageJ software (National Institutes of Health). A list of PCR primer sequences is available in Supplementary Table S1.

### Subcellular fractionation

Nuclei from Flag-tagged DPF2-S or DPF2-L expressing mouse ESCs were purified as described (Damianov et al., 2016; Grabowski, 2005) and were lysed for ∼5 min in five volumes of ice-cold lysis buffer (20 mM HEPES-KOH (pH 7.5), 150 mM NaCl, 1.5 mM MgCl2, 0.6% Triton X-100, and protease inhibitor cocktail). Soluble nucleoplasmic (NP) and high molecular weight (HMW) chromatin fractions were separated by centrifugation at 20,000 x g for 5 min at 4°C. To extract protein complexes, an equal volume of lysis buffer was added to the HMW pellet. Soluble NP and HMW fractions were incubated at room temperature on a rotator with 5 U/ml of Benzonase nuclease (Sigma) until the HMW pellet was resuspended. Both the NP and HMW fractions were then cleared by centrifugation at 20,000 x g for 10 min at 4°C for immunoprecipitation experiments.

### Immunoprecipitation of BAF complex

Soluble nucleoplasmic (NP) and the high molecular weight (HMW) fractions were incubated overnight at 4°C with 20 ul packed M2 FLAG agarose beads (Sigma). Beads were washed four times with IP wash buffer (20 mM HEPES-KOH (pH 7.5), 150 mM NaCl, 1.5 mM MgCl2, 0.05% Triton X-100, and protease inhibitor cocktail). Flag-tagged proteins were eluted from the beads using 150 ng/ml of 3xFLAG peptide (Sigma) in 50–100 ul of elution buffer (20 mM HEPES-KOH (pH 7.5), 100 mM NaCl, 1.5 mM MgCl2, and protease inhibitor cocktail) for 1 hr at 4°C in a Thermomixer. Eluted proteins/complex were subject to SDS-PAGE electrophoresis followed by protein staining (SYPRO Ruby Protein Gel Stain, Thermo Fisher Scientific) and immunoblotting.

### Immunoblotting

Immunoblotting was performed on total cell lysates using RIPA buffer (Boston BioProducts) supplemented with protease inhibitors (Roche), or on the soluble nucleoplasmic (NP) and high molecular weight (HMW) fractions of the nuclei. Lysates were diluted in 5X SDS loading buffer (50 mM Tris-Cl (pH 6.8), 0.05% bromophenol blue, 2% SDS, 10% glycerol, and 0.1 M DTT), heated at 95 C° for 5 minutes, loaded onto 12% polyacrylamide SDS PAGE gels and run under standard electrophoresis conditions. Transfer was performed on a Trans-Blot SD Semi-Dry Transfer Cell (Bio-Rad) onto Immobilon-FL PVDF membranes (EMD Millipore). The membranes were incubated with primary antibodies overnight at 4 °C. Primary antibodies used in this study are anti-DPF2 (1:500, Abcam, #ab134942), anti-Flag M2 (1:2000, Sigma Aldrich, #F3165), anti-BAF60a (SMARCD1) (1:200, Santa Cruz Biotechnology, #sc-135843), anti-Ini1 (SMARCB1) (1:150, Santa Cruz Biotechnology, #sc-166165), and anti-Histone H3 (1:1000, Abcam, #ab1791-100). For fluorescent detection of primary antibody, the membranes were probed with ECL Plex Cy3- or Cy5-conjugated goat anti-mouse or anti-rabbit secondary antibodies (Cytiva) and scanned on an Amersham Typhoon 5 Phosphorimager (GE Healthcare).

### RNA sequencing and data analysis

To identify gene expression changes during neuronal differentiation, poly(A)-plus RNA was isolated from the Hb9-ESCs, ESC-derived NPCs (DIV-21), and ESC-derived GNs (DIV-10). Paired-end strand specific libraries were constructed using the TruSeq stranded mRNA (polyA) Librabry Prep Kit (Illumina) and were subject to 100 nt paired-end sequencing (Illumina NovaSeq S1 (XP)) to generate 22∼30 million mapped reads per sample at the UCLA Neuroscience Genomics Core (UNGC) facility. RNA-seq reads were aligned to the mouse NCBI38/mm10 reference genome using STAR (Dobin et al., 2013). We used FeatureCounts (Liao et al., 2014) to generate counts and DESeq2 (Love et al., 2014) was used to identify differentially expressed genes with a 1.5 fold change and a p-value cutoff of < 0.05. Gene expression (TPM) was calculated using Kallisto (Bray et al., 2016). Gene Ontology analyses were performed using Metascape (Zhou et al., 2019).

### Chromatin immunoprecipitation coupled with high-throughput sequencing (ChIP-seq)

ChIP-seq was performed as previously described (Barish et al., 2010; Purbey et al., 2017) using anti-DPF2 antibody (Abcam, #ab134942). Briefly, approximately 20 million embryonic stem cells or neural progenitor cells were used per sample. Cells were then treated with 2mM DSG followed by treatment with 1% Formaldehyde. After cross-linking, cells were sonicated on a Covaris M220-focused ultrasonicator. We then checked the distribution of chromatin using DNA electrophoresis to ensure that chromatin shearing generated DNA fragments between 200-500 bps. ChIP-seq libraries were prepared using the Kapa Hyper Prep Kit (Kapa Biosystems), followed by sequencing (Illumina HiSeq 3000).

### ChIP-seq data analysis

Trimmed sequences were aligned to the mouse NCBI38/mm10 reference genome using STAR (Dobin et al., 2013). PCR duplicated reads were filtered using SAMtools (Li et al., 2009). HOMER (Heinz et al., 2010) was used to call peaks, creating bedgraphs, and finding motifs.

BedTools (Quinlan and Hall, 2010) was used to merge peaks. Peaks were quantitated across samples and normalized to million reads per sample and peak length (RPKM) using SeqMonk (https://www.bioinformatics.babraham.ac.uk/projects/seqmonk/). Differential binding peaks were selected using DEseq2 (Love et al., 2014). A *P*-value cutoff of <0.05 was used to identify differential peaks. Peaks were annotated to genes using HOMER (Heinz et al., 2010). Deeptools2 (Ramírez et al., 2016) was used to generate the metaplots to show the distribution of chromatin marks around or within the differential ChIP-seq peaks.

Public ChIP-seq data sets were obtained from previous studies (Beagan et al., 2017; Chronis et al., 2017; Kubo et al., 2021; Mateo et al., 2015; Nishi et al., 2015; Oldfield et al., 2014; Wapinski et al., 2013).

### Assay for transposase-accessible chromatin with sequencing (ATAC-Seq)

ATAC-seq was performed as previously described (Buenrostro et al., 2015). Briefly, about 50,000 cells were harvested and treated with 500 ul of cold lysis buffer (10 mM Tris-HCl (pH 7.4), 10 mM NaCl, 3 mM MgCl2, 0.1% IGEPAL CA-630). Lysed cells were then treated with 2xTD reaction buffer (Illumina, #FC-121-1030) and TDE1 Nextera Tn5 Transposase (Illumina, #FC-121-1030) and incubated for 30 minutes at 37°C. Transposed DNA was then purified using Qiagen MinElute PCR purification kit and eluted with elution buffer (10 mM Tris buffer, pH 8). ATAC-seq libraries were then prepared using Custom Nextera PCR primers and NEBNext High-Fidelity 2x PCR Master Mix (NEB, #M0541).

### ATAC-seq data analysis

Sequences were trimmed using Cutadapt (Martin, 2011) and trimmed reads were aligned to the mm10 using STAR (Dobin et al., 2013). PCR duplicated reads were filtered using SAMtools (Li et al., 2009). ATAC-seq peaks are called using MACS3 (Zhang et al., 2008) and merged peaks were created using BedTools (Quinlan and Hall, 2010). Peaks were quantitated across samples and normalized to million reads per sample and peak length (RPKM) using SeqMonk (https://www.bioinformatics.babraham.ac.uk/projects/seqmonk/). Differentially accessible peaks were determined using DEseq2 (Love et al., 2014).

### Chromatin state enrichment assay

Chromatin state enrichment analyses were done as previously described (Ernst and Kellis, 2017). We used a 18-state ChromHMM model for mouse ESCs as described before (Chronis et al., 2017). For NPCs, we used ChromHMM annotations from a 15-state model for the forebrain specific 15-state ChromHMM model at developmental day E15.5, a stage where most cells are in a NPC-like state (Gorkin et al., 2020). For the analysis of binding in binding in both ESCs and NPCs, we also used a previously described universal 100-state stacked ChromHMM model (Vu and Ernst, 2022).

## SUPPLEMENTARY FIGURES

**Figure S1.**
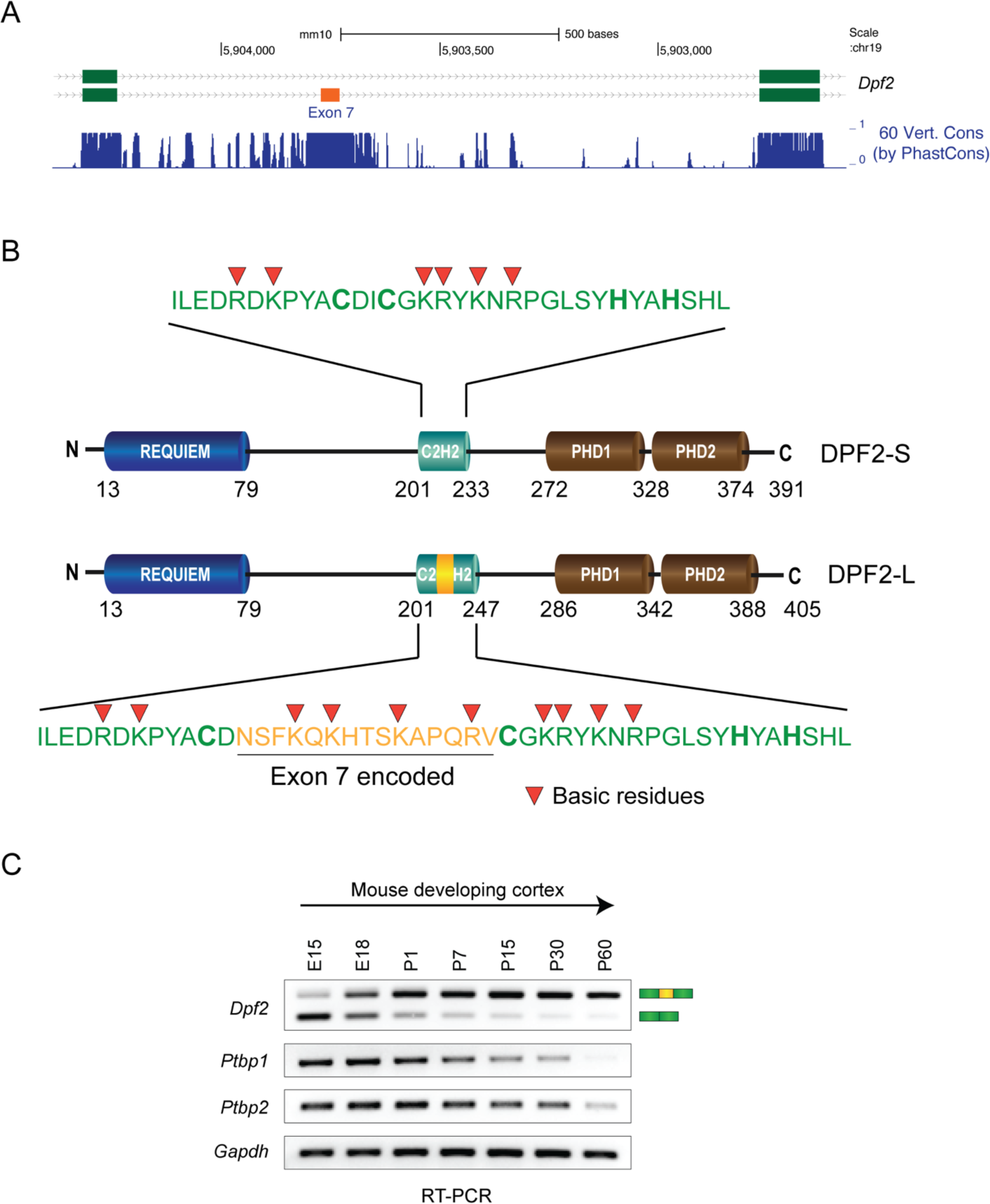
Developmental splicing switch of highly conserved *Dpf2* exon 7. Related to Figure 1. **(A)** Genome browser track of *Dpf2* exon 7 showing 60 vertebrate conservation by PhastCons. **(B)** Schematic of the canonical DPF2-S and the longer DPF2-L isoforms with amino acid sequences of the C2H2-type zinc finger domain where exon 7 inclusion incorporates 14 amino acids (shown in orange). Amino acid numbers are shown, and basic residues are indicated by red arrowheads. **(C)** RT-PCR assays of *Dpf2* exon 7 alternative splicing in mouse cortices at embryonic days E15 and E18, and postnatal days P1, P7, P15, P30, and P60. Expression *Ptbp1* and *Ptbp2* RNA is shown below with *Gapdh* as a loading control.

**Figure S2.**
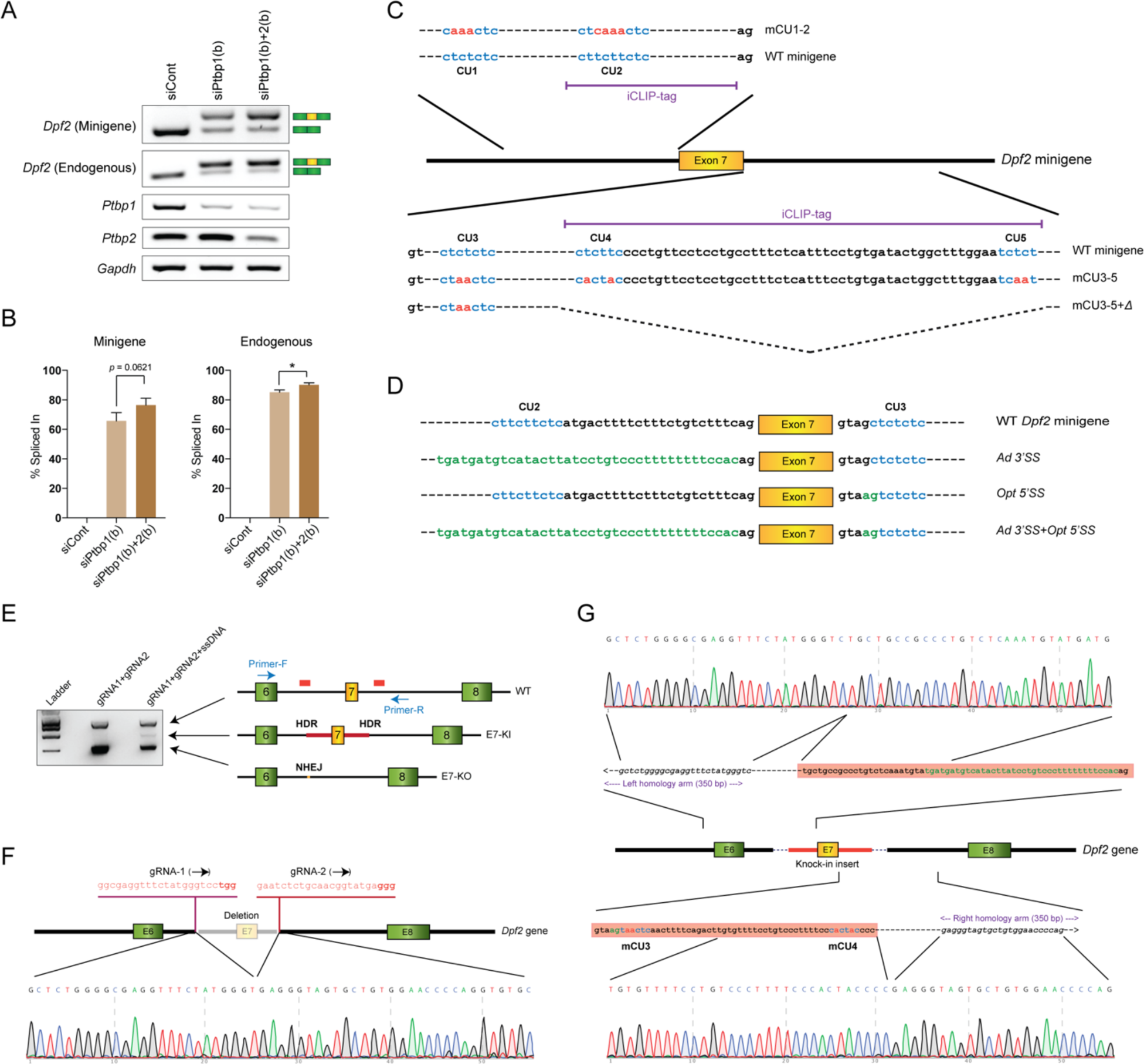
PTBP1-mediated regulation of *Dpf2* exon 7 alternative splicing and establishment of genome-edited mouse ESC lines. Related to Figures 2 & 3. **(A)** RT-PCR assay of exon 7 splicing in the *Dpf2* minigene and endogenous *Dpf2* transcripts after siRNA knockdown of *Ptbp1* and *Ptbp2* by a second set of siRNAs. Expression of *Ptbp1*, *Ptbp2*, and *Gapdh* RNA are shown below. **(B)** Bar graphs of quantified RT-PCR results calculated as exon 7 percent spliced in (PSI) for the *Dpf2* minigene and endogenous *Dpf2*. Each bar represents the mean value (+/-SD) of three independent biological replicates. *: p < 0.05 (Student’s t-test). **(C)** Schematic of the introns flanking *Dpf2* exon 7 containing the CU-rich elements. Mutations disrupting the CU-rich motifs are shown in red. Dotted lines in mutant mCU3-5+Δ indicate the deleted region containing a PTBP1 iCLIP cluster. **(D)** Sequences of the 3’ and 5’ splice sites flanking exon 7. Mutations that replace the 3’ splice site or convert the weak 5’ splice site to an optimal site are shown in green. **(E)** PCR genotyping of pooled CRISPR/Cas9-mediated genome edited mouse ESCs by PCR. Schematic on the right shows the origin of each PCR band. **(F)** Sanger sequencing results confirming deletion of the 438 bp region harboring exon 7 and its flanking intronic sequences in the E7-KO clones. The positions of the gRNA target sequences, and their orientation are indicated. The PAM motifs are shown in bold red. **(G)** Sanger sequencing confirming a 161 bp knock-in insert harboring exon 7 and flanking intronic segments carrying an optimal 5’ splice site (green), an adenovirus 3’ splice site (green), and mutated CU-rich elements (red).

**Figure S3.**
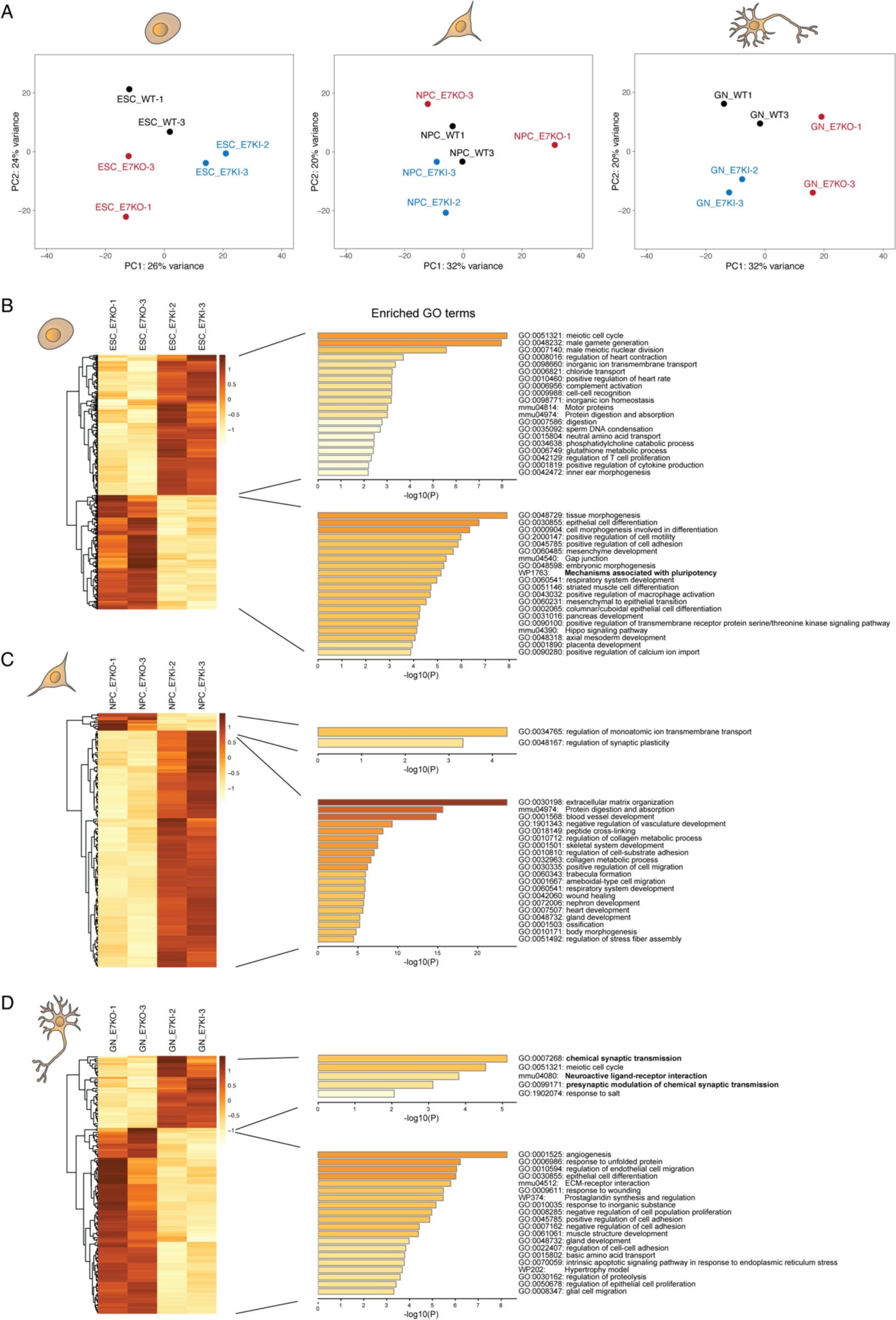
Alternative DPF2 isoforms regulate distinct programs of gene expression during neuronal differentiation of ESCs. Related to Figure 4. **(A)** Principal component analysis (PCA) of RNA-seq samples from ESCs, ESC-derived NPCs, and ESC-derived GNs. **(B-D)** Heatmaps showing differentially expressed genes (DEGs) detected by RNA-seq in mouse ESCs (B), NPCs (C), and GNs (D) expressing either DPF2-S or DPF2-L. Metascape analysis of enriched Gene Ontology (GO) terms for biological processes associated with the DEGs are shown on the right.

**Figure S4.**
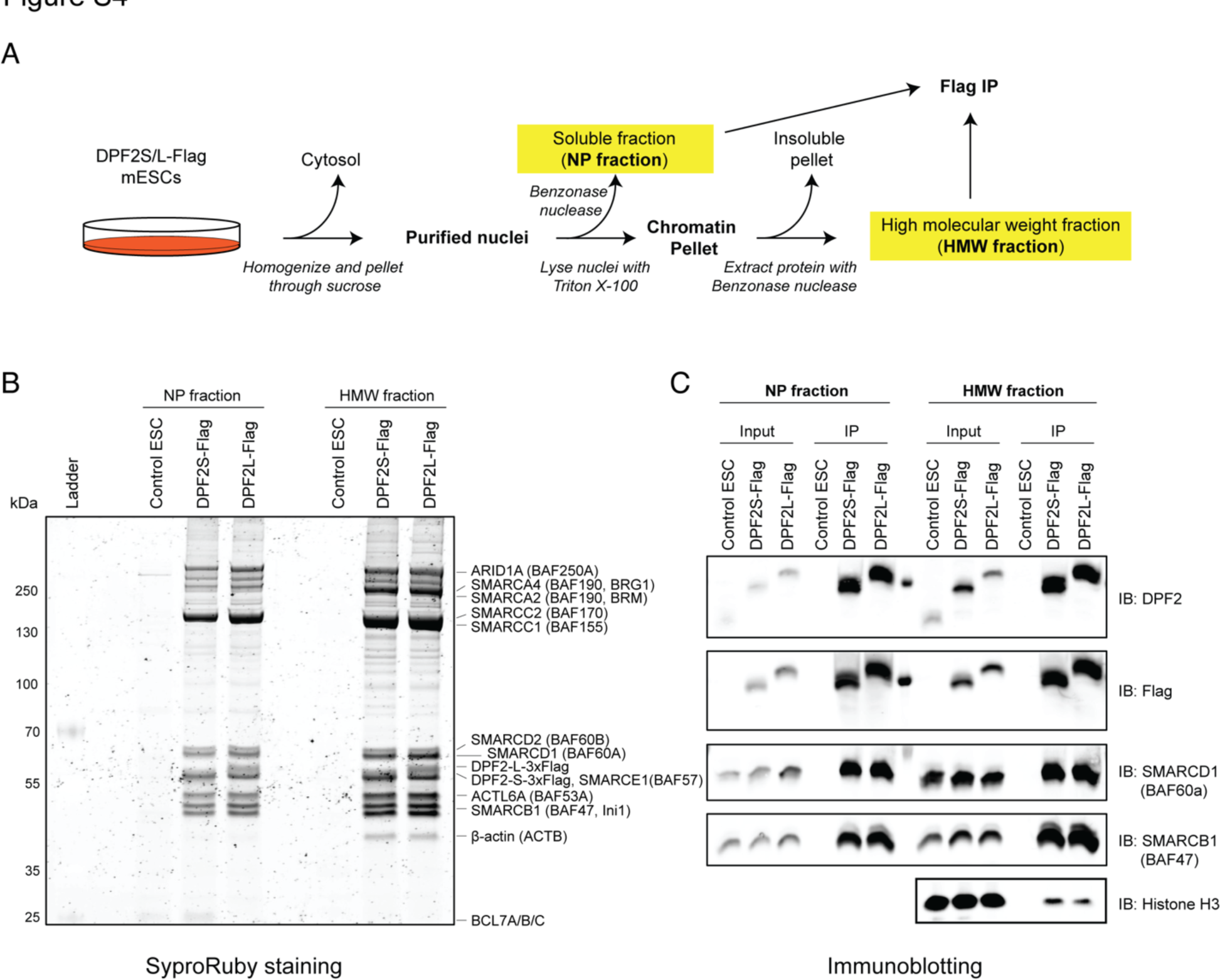
Purification of BAF complex with Flag-tagged DPF2-S or DPF2-L. Related to Figure 5. **(A)** Schematic showing the method of extracting soluble nucleoplasmic (NP) and high molecular weight fractions (HMW) from the nuclei. **(B)** Protein staining (SyproRuby) of the Flag peptide eluates following anti-Flag immunoprecipitation from the NP and HMW fractions. Proteins corresponding to each band are labeled as identified by mass spectrometry and immunoblot. **(C)** Immunoblot analyses of BAF complex subunits in the IP eluates from the NP and HMW fractions.

**Figure S5.**
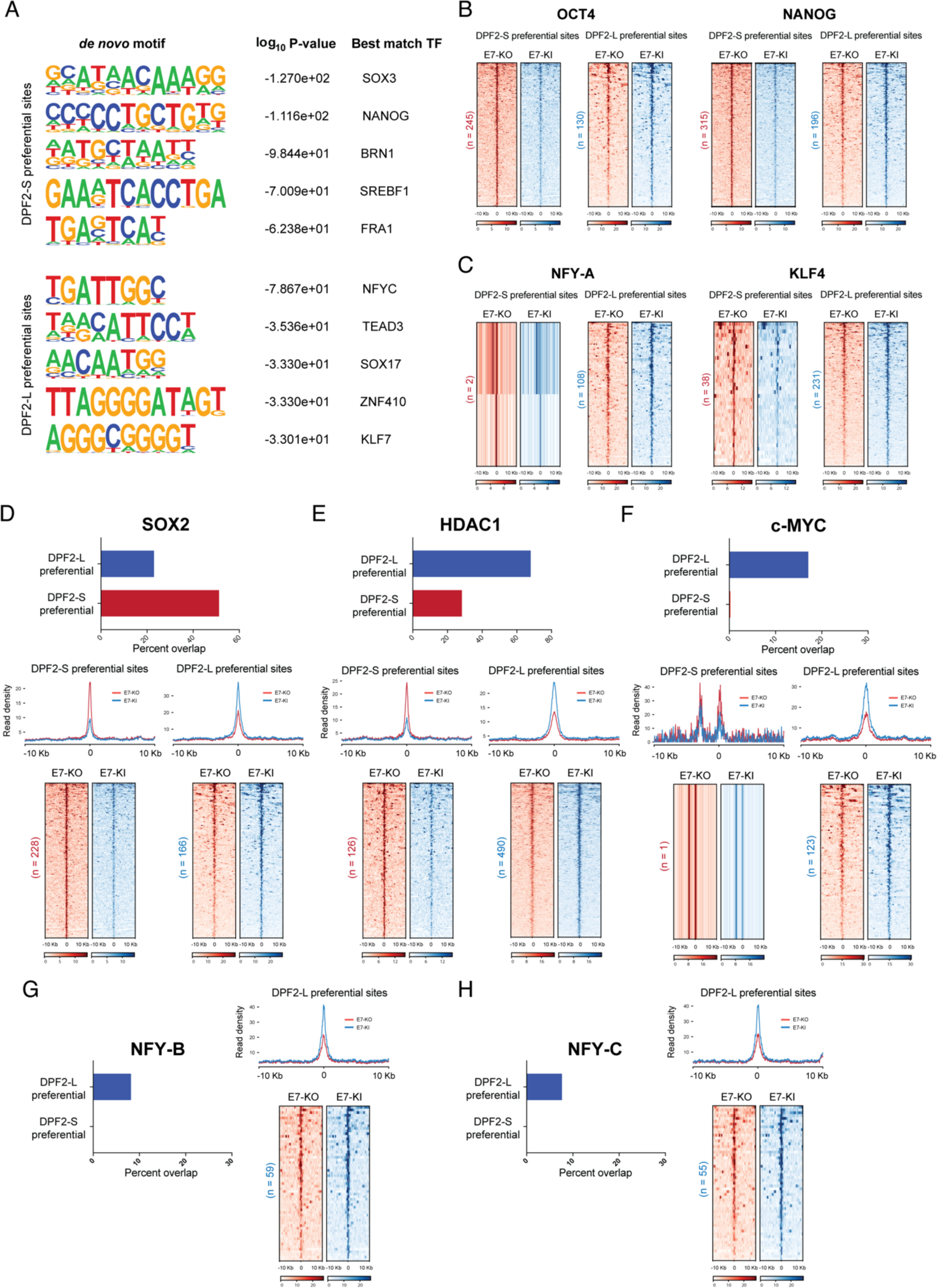
Transcription factor binding at DPF2-S or DPF2-L preferential binding sites in mouse ESCs. Related to Figure 5. **(A)** Enrichment of *de novo* motifs in the DPF2-S and DPF2-L preferential binding sites in ESCs determined by HOMER. Log_10_ *P-*values and best match transcription factors are listed. **(B-C)** Normalized tag density profiles of average signal intensities for DPF2-S and DPF2-L preferential binding sites overlapping with known OCT4, NANOG (B), NFY-A, and KLF4 (C) binding sites determined by ChIP-seq (Chronis et al., 2017; Oldfield et al., 2014). **(D-H)** Percent overlap and metaplots of average signal intensities of DPF2-S and DPF2-L preferential binding sites with known SOX2 (D), HDAC1 (E), c-MYC (F), NFY-B (G), and NFY-C (H) binding sites in ESCs (Chronis et al., 2017; Oldfield et al., 2014). Bottom panel shows metaplots and normalized tag density profiles of average signal intensities of DPF2-S and DPF2-L preferential binding sites overlapping with the transcription factor binding sites indicated above.

**Figure S6.**
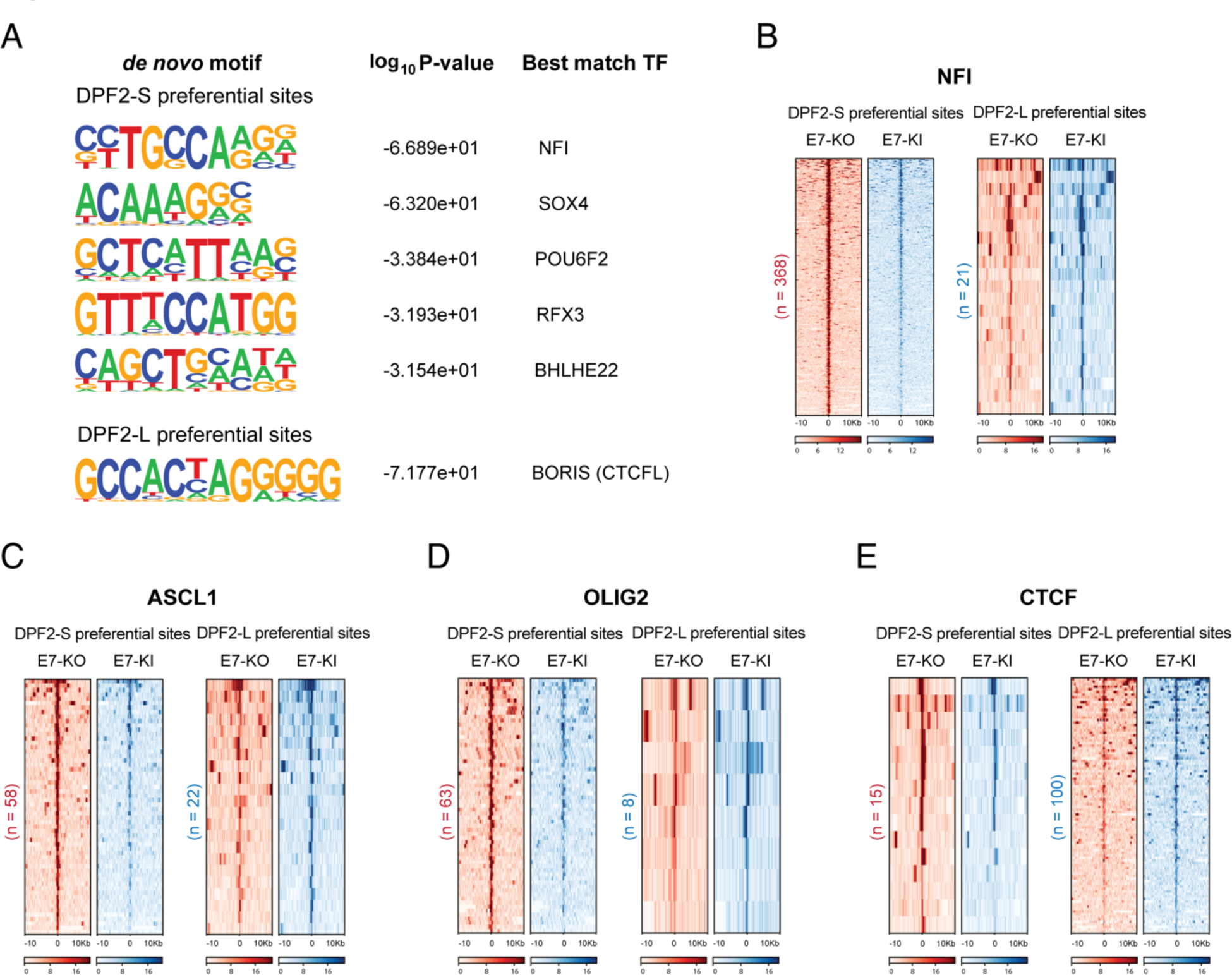
Transcription factor binding at DPF2-S or DPF2-L preferential binding sites in mouse NPCs. Related to Figure 6. **(A)** Enrichment of *de novo* motifs determined by HOMER analysis of the DPF2-S and DPF2-L preferential binding sites in NPCs. Log_10_ *P-*values and best match transcription factors are listed. **(B-E)** Normalized tag density profiles of average signal intensities for the DPF2-S and DPF2-L preferential binding sites overlapping with known NFI (B), ASCL1 (C), OLIG2 (D), and CTCF in NPC’s (E) (Beagan et al., 2017; Mateo et al., 2015; Nishi et al., 2015; Wapinski et al., 2013).

**Figure S7.**
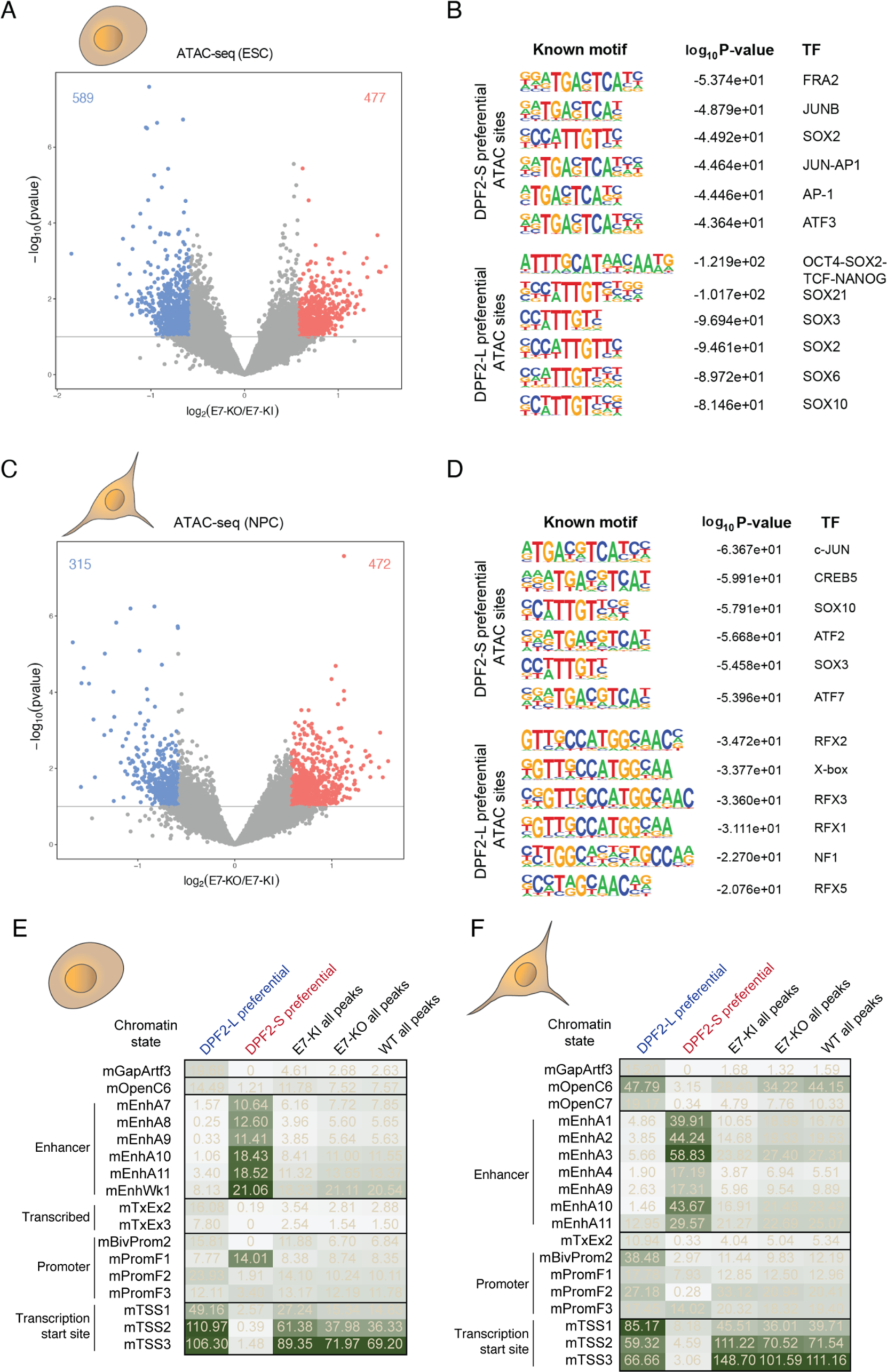
Preferential open chromatin sites and chromatin state enrichments in ESC and NPC. Related to Figures 5, 6, & 7. **(A)** Volcano plot of sites exhibiting differential chromatin accessibility detected by ATAC-seq in mouse ESCs expressing either DPF2-S or DPF2-L. Significantly different sites exhibiting greater than 1.5-fold changes (*P*-value < 0.05) between DPF2-S and DPF2-L are labeled as red and blue dots, respectively. **(B)** Motif enrichment analysis (HOMER) for known transcription factor motifs within DPF2-S and DPF2-L preferential open chromatin sites in ESCs. Corresponding *P-*values and transcription factors are listed for each motif. **(C)** Volcano plot of sites exhibiting differential chromatin accessibility detected by ATAC-seq in NPCs. Significantly different (*P*-value < 0.05) DPF2-S and DPF2-L preferential open chromatin sites are labeled as red and blue dots, respectively. **(D)** Motif enrichment analysis (HOMER) for known transcription factor motifs within DPF2-S and DPF2-L preferential open chromatin sites in NPCs. Corresponding *P-*values and transcription factors are listed. **(E-F)** Chromatin state enrichment analysis of DPF2-S and DPF2-L preferential binding sites in ESCs (D) and ESC-derived NPCs (E) using the universal chromatin state annotations from a stacked ChromHMM model (Vu and Ernst, 2022). The rows represent different chromatin states, and their state mnemonics are displayed. Columns show fold enrichments (ChromHMM emission probabilities) of each state within DPF2-S and DPF2-L preferential binding sites (columns 1 and 2), as well as in other peak sets, colored from highest to lowest intensities. A subset of states showing the largest changes in fold enrichments between DPF2-S and DPF2-L preferential binding sites are shown in this figure. Enrichments for the complete set of 100 states is shown in Supplementary Table S6.

**Supplementary Table S1.**
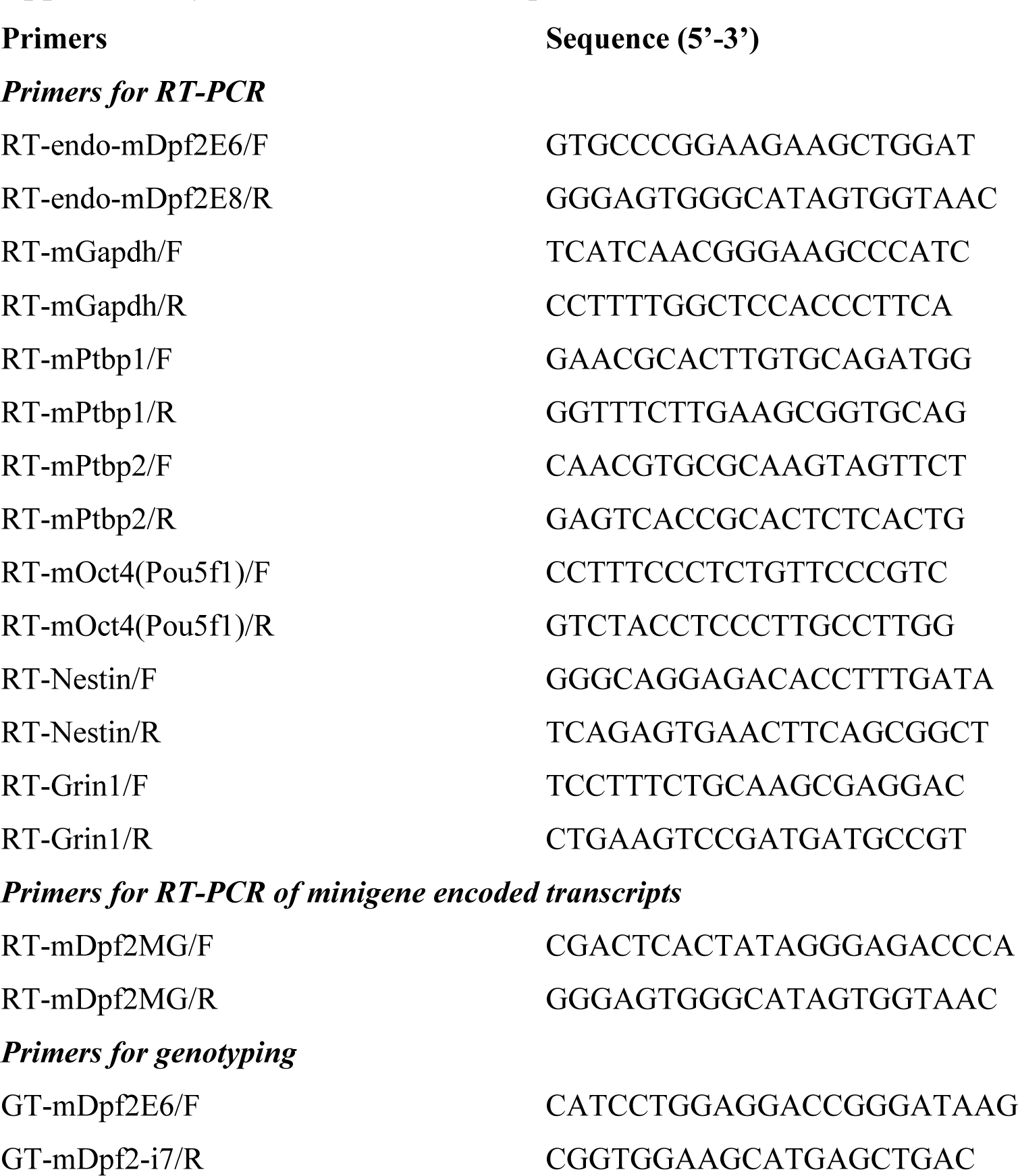
List of PCR primers.

**Supplementary Table S2.** List of siRNAs.

| **siRNA** | **Name** | **Catalog number** |
| --- | --- | --- |
| siNTC | Silencer Select Negative Control #1 | Thermo Fisher Scientific, 4390843 |
| siPtpb1 (1) | Silencer Select siRNAs against *Ptpb1* | Thermo Fisher Scientific, s72335 |
| siPtpb1 (2) | Silencer Select siRNAs against *Ptpb1* | Thermo Fisher Scientific, s72337 |
| siPtpb2 (1) | Silencer Select siRNAs against *Ptpb2* | Thermo Fisher Scientific, s80148 |
| siPtpb2 (2) | Silencer Select siRNAs against *Ptpb2* | Thermo Fisher Scientific, s80149 |

**Supplementary Table S3.** List of antibodies.

| **Name** | **Manufacturer** | **Catalog number** |
| --- | --- | --- |
| Anti-DPF2 | Abcam | ab134942 |
| Anti-Flag M2 | Sigma Aldrich | F3165 |
| Anti-BAF60a | Santa Cruz Biotechnology | sc-135843 |
| Anti-Ini1 | Santa Cruz Biotechnology | sc-166165 |
| Anti-Histone H3 | Abcam | ab1791-100 |

## Notes

### Competing Interest Statement

The authors have declared no competing interest.

